# Buildup and bistability in auditory streaming as an evidence accumulation process with saturation

**DOI:** 10.1101/2020.01.24.917799

**Authors:** Quynh-Anh Nguyen, John Rinzel, Rodica Curtu

**Affiliations:** Department of Mathematics, The University of Iowa, Iowa City, Iowa, United States of America; Center for Neural Science, New York University, New York, New York, United States of America; Courant Institute of Mathematical Sciences, New York University, New York, New York, United States of America; Iowa Neuroscience Institute, Human Brain Research Laboratory, Iowa City, Iowa, United States of America

## Abstract

A repeating triplet-sequence *ABA*_ of non-overlapping brief tones, *A* and *B*, is a valued paradigm for studying auditory stream formation and the cocktail party problem. The stimulus is “heard” either as a galloping pattern (integration) or as two interleaved streams (segregation); the initial percept is typically integration then followed by spontaneous alternations between segregation and integration, each being dominant for a few seconds. The probability of segregation grows over seconds, from near-zero to a steady value, defining the buildup function, BUF. Its stationary level increases with the difference in tone frequencies, *DF*, and the BUF rises faster. Percept durations have *DF* -dependent means and are gamma-like distributed. Behavioral and computational studies usually characterize triplet streaming either during alternations or during buildup. Here, our experimental design and modeling encompass both. We propose a pseudo-neuromechanistic model that incorporates spiking activity in primary auditory cortex, A1, as input and resolves perception along two network-layers downstream of A1. Our model is straightforward and intuitive. It describes the noisy accumulation of evidence against the current percept which generates switches when reaching a threshold. Accumulation can saturate either above or below threshold; if below, the switching dynamics resemble noise-induced transitions from an attractor state. Our model accounts quantitatively for three key features of data: the BUFs, mean durations, and normalized dominance duration distributions, at various *DF* values. It describes perceptual alternations without competition per se, and underscores that treating triplets in the sequence independently and averaging across trials, as implemented in earlier widely cited studies, is inadequate.

**Author summary:** Segregation of auditory objects (auditory streaming) is widely studied using ambiguous stimuli. A sequence of repeating triplets *ABA*_ of non-overlapping brief pure tones, *A* and *B*, frequency-separated, is a valued stimulus. Studies typically focus on one of two behavioral phases: the early (say, ten seconds) buildup of segregation from the default integration or later spontaneous alternations (bistability) between seconds-long integration and segregation percepts. Our experiments and modeling encompass both. Our novel, data-driven, evidence-accumulation model accounts for key features of the observations, taking as input recorded spiking activity from primary auditory cortex (as opposed to most existing, more abstract, models). Our results underscore that assessing individual triplets independently and averaging across trials, as in some earlier studies, is inadequate (lacking neuronal-accountability for percept duration statistics, the underlying basis of buildup). Further, we identify fresh parallels between evidence accumulation and competition as potential dynamic processes for choice in the brain.

## Introduction

Stimulus sequences of interleaved *A* and *B* pure tones have been widely used in studying segregation of distinct objects in an auditory scene (auditory streaming), in human psychophysics [1–6], invasive neurophysiology [1, 3, 7, 8], or in experiments implementing both [9]. A valued stimulus is triplet-streaming *ABA*_ with the tone frequency difference, *DF*, as a tunable parameter [10]; Fig 1. For small *DF* human listeners most likely perceive integration (one galloping rhythm); for *DF* large, segregation dominates (two simultaneously heard parallel streams). The initial percept is typically integration but within seconds the probability of segregation increases (“the buildup phase”) and perceptual switching eventually occurs (“perceptual bistability”). Alternating percepts have variable durations, described by either gamma or lognormal distributions [2]. Time courses of spiking activity (macaque, primary auditory cortex, A1, [1]) show dynamical features (adaptation over 1-2 seconds) that were interpreted as neural correlates of buildup, although the behavioral and physiological experiments were not conducted together [1, 3].

**Fig 1.**
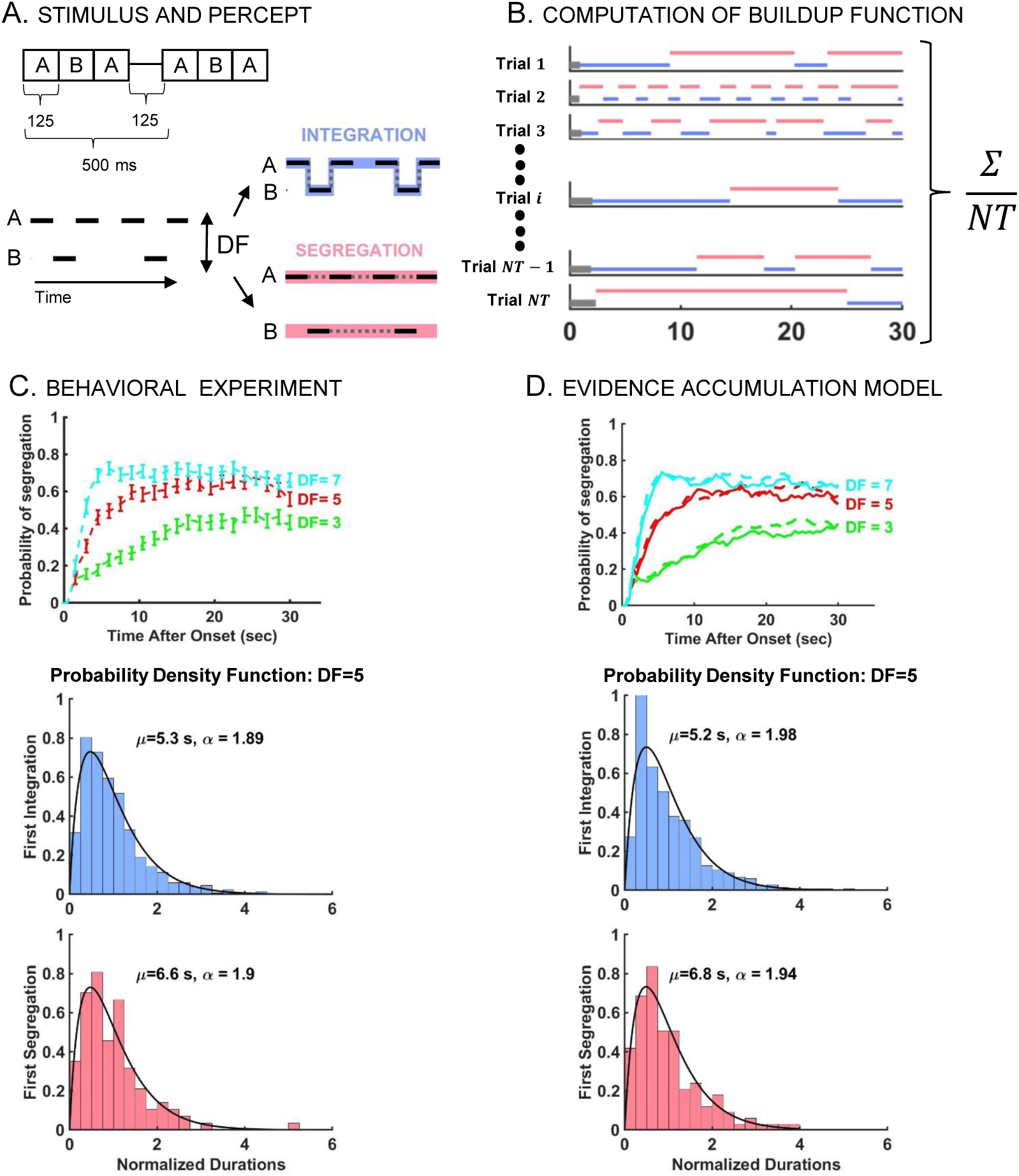
Stimulus, buildup, and distribution of first percept durations in auditory streaming of triplets. A: Stimulus paradigm (left) used for behavioral experiments and corresponding percept types (right). Stimulus consists of sequences of high (*A*) and low (*B*) pure tones presented as repeated triplets *ABA*_ where ‘_’ denotes silent gap. Depending on *DF* between tones *A* and *B*, there are two fundamental percepts: integration (*I*; blue), one connected stream with galloping rhythm, and segregation (*S*; red), two parallel streams of high tone *A*_*A*_*A*_*A*_ and low tone _*B*___*B*__ occurring simultaneously. B: Computation of the buildup function (time course of probability of *S*) obtained by determining the frequency of occurrence of *S* over all trials at each time point *τ* up to 30 s (45 trials for each *DF* and subject; 15 subjects). Non-*S* includes both latency (gray) and *I* (blue) states. Due to latency, the buildup function always starts at 0 even though the first percept is not necessarily *I*. For example, in Trial 2, the first percept is *S*. C: Experimental-based psychometric buildup function (upper panel) and distribution of first percept durations (middle and lower panels). Buildup functions are computed for *DF* =3 (green), 5 (red), and 7 (cyan). The error bars indicate 95% CI around the mean using statistical bootstrapping. Durations are normalized by dividing by mean duration. Likelihood ratio test confirms that normalized first percept durations are gamma distributed – shown here at *DF* =5 for *I* (*N* =533, *p*=0.49) and *S* (*N* =114, *p*=0.47). The shape parameters *α*, obtained by Maximum Likelihood Estimation (MLE), and the mean durations *µ* are indicated in the graphic. D: Model-based simulated buildup function (upper panel) and distribution of normalized first percept durations (middle and lower panels). Buildup functions from the evidence accumulation model (EVA; solid) closely resemble those from the behavioral experiment (dashed, also in C). Normalized first percept durations are gamma distributed (shown at *DF* =5). Similar results are obtained for other *DF* values; see S1 Fig.

Dynamics of buildup and/or perceptual alternation for ambiguous auditory stimuli were described by computational models based on signal processing [1, 3, 11–13], competition dynamics [6, 14], coupled-oscillator patterning [15, 16], evidence accumulation [5], and statistical descriptions [17]; also reviews by [18] and [19]. However, with few exceptions (e.g. [1, 6]) these models did not incorporate neurophysiological data. Furthermore, experimental and modeling studies primarily focused on either buildup, describing the probability of segregation during short, tens of seconds, trials [1, 3, 4, 20, 21], or on the stationary phase of alternations, characterizing the statistics of percept durations over long, several minutes, trials [2, 5, 6].

Here we designed the experiment (30 s trials with many trials per condition/subject) so that we could characterize these features simultaneously. Then we proposed a model that takes spike-recordings from A1 as input, and accounts for both the behavioral time course of buildup and the observed duration statistics during alternations, over a range of *DF* values: 3, 5, 7 semitones. Our model is neuromechanistic-like, transforming the neuronal input for processing in two evidence-accumulators downstream of A1. From the input-sensory level, sampling of spike counts across A1-units provides a measure for the contribution of each triplet to the evidence-accumulation stage; if evidence against the current percept exceeds a threshold then a perceptual switch occurs and accumulation resets. This approach parallels in spirit Barniv and Nelken’s model [5] although that was implemented from a Bayesian-viewpoint. Our model is data-driven: input is neural-based; initial parameters are estimated from our behavior data (mean probability of segregation) then fine-tuned to match the gamma-distributed percept durations.

We propose that although the model is not competition-based it shares some features of such approaches: Adaptation is key in competition dynamics; evidence accumulation might be viewed as recovery from adaptation. Matching duration statistics with competition requires some balancing of noise and adaptation [22]; its analogue is the interplay between accumulation and noise. Adaptation strength, when set near the boundary between noise-free oscillatory and noise-driven attractor dynamics, constrains dominance durations [23]; comparably, our accumulators have a novel feature of saturation which if set below but near the switching threshold, produces observed statistics only if adequate noise is present.

Importantly, our modeling highlights that accounting for the duration statistics of behavioral data is key when studying auditory bistable perception. With quantitative matches to these data the buildup phase is then naturally reproducible by an alternating renewal process [17]. We show that a widely cited signal-detection approach [1, 3, 12, 21], based on treating each triplet independently without accumulation, that overlooked this crucial feature does not account for the single-trial percept duration statistics. We argue for caution when applying it to test neural-inspired behavioral hypotheses.

## Results

We first outline the rationale for our study and presentation. In behavioral experiments human participants continuously reported their ongoing perception, integration or segregation, which after analysis yielded distributions for percept durations (Section A). We introduce the essence of our **EV**idence **A**ccumulation (EVA) model in a basic form (Section B.1). With each triplet we suppose there is an incremental urge *r*, to switch from the current percept/interpretation to the alternate one; *r* is the “drift” rate for the event sequence that, with zero-mean noise, drives fluctuating accumulation in the EVA model that eventually surpasses threshold. We illustrate that this basic model captures the duration statistics for a chosen case, near-equidominance. We next elaborate the model by formulating a neuronal basis for evaluating *r* (Section B.2). We utilize the single-unit spike counts for *A*-tone selective A1 neurons recorded over a range of experimental conditions [1] and applied them to our case of *DF* =3, 5, 7. The relative responses to *B*-tones are viewed, according to the population separation hypothesis [7], as evidence for segregation (against integration) when spike counts are generally smaller, or against segregation when larger. A challenge arises. If *N*_*in*_ A1 neurons are recorded the spike count deviates from the mean like 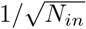. Thus, if *N*_*in*_ is large and one supposes a fixed threshold for signal detection, the classification based on spike counts becomes binary and problematic for resolving a perceptual response that is graded over conditions. Our full EVA model (Section B.3) attempts to meet the challenge by having a two-layer pre-processing stage that includes *N*_*sl*_ units, each of them sampling a few A1 neurons (*N*_*in*_ not large). The proportion of *N*_*sl*_ units which respond to thresholded activity over neuronal ensembles in A1 provides the incremental evidence, the value of *r*, for the accumulator that favors integration. The complementary proportion of *N*_*sl*_ units that do not respond to thresholded A1 activity provides the incremental evidence to the accumulator against integration.

Our approach overcomes two shortcomings of a well-known signal detection model for auditory streaming [1]. The Micheyl et al treatment [1] does not account for single-trial data, the duration distributions which form the basis for computing the buildup function (BUF); it averages across trials, without accumulating evidence event-to-event. The signal detection scheme of Micheyl does not resolve, with *N*_*in*_ large, a family of BUFs that show gradation across conditions.

In short, we combined neural data from [1] with behavioral data from our experiments (see Section A) to investigate if the signal detection model when applied on a single-trial basis could yield percept durations in a self-consistent fashion. We found it did not; and moreover that it was unable to fit buildup functions that, for different stimulus conditions, were graded, not widely separated (Section C). We then developed a neural-based evidence accumulation-like explanation of the observed data, as alternative to explicit competition, and with the advantage of being intuitive (Section B).

### A. Auditory triplet-streaming

#### A.1. Experimental protocol

Fifteen human subjects with normal hearing listened to sequences of repeating *ABA*_ triplets and were instructed to continuously report their ongoing percept by selectively pressing one of two different buttons on a keypad. Subjects began reporting their percept typically 2 s after stimulus onset as integration (*I*; a single, coherent stream *ABA*_*ABA*_) or segregation (*S*; two distinct streams *A*_*A*_*A*_*A*_ and _*B*___*B*_). Stimuli were sequences of triplets *ABA*_ that consisted of alternating high (*A*) and low (*B*) pure tones followed by a 125 ms silent pause “_” (Fig 1A-B). In total, triplets were 500-ms in duration and were repeated 60 times per trial. Tones were separated in frequency by *DF* semitones chosen from three conditions (*DF* = 3, 5, 7) with each condition being presented five times per experimental block (nine blocks total). This resulted in group data (from 15 subjects and 45 trials per subject) with 675 30-s trials for each of three *DF* values.

#### A.2. Behavioral task performance

For each *DF* condition the buildup function was constructed by computing the probability of segregation from trial-averaging (Fig 1 B-C). The buildup functions started at zero and increased over time before stabilizing to certain *DF* -dependent asymptotic values, similar to reports by [1, 3, 5, 12]. They started at zero due to the latency period (when no percept was identified) and not because the initial percept was *I*; see *Methods*, also [4]. While *I* first percepts were indeed more likely, *S* first percepts were reported too. The proportion of segregation as initial percept increased with *DF* from 103 out of 675 trials at *DF* =3 to 137 at *DF* =5 and 220 at *DF* =7. The probability of segregation increased faster and reached higher levels at larger *DF*, with transient times of approximately 16, 10, 5 s after stimulus onset and with asymptotic values 0.45, 0.6, and 0.65 at *DF* = 3, 5, 7 respectively.

We computed distributions of normalized phase durations for subsequent durations, separately for each *DF*, and found them to be gamma-like, consistent with previous results on subsequent percepts [2, 5, 6]. Herein we report that duration distributions of the first percept are also gamma-like (Fig 1C; see also S1 Fig). We used statistical bootstrapping to compute the shape parameter *α* of each gamma distribution (see *Methods*), and determined that *α* ≈ 2 for normalized first durations and *α* ≈ 2.6 for subsequent durations. The distributions satisfied the scaling property *γ*_1_ ≈ 2*CV* with skewness *γ*_1_ and coefficient of variation *CV* ≈ 0.7 and *CV* ≈ 0.6 respectively, similar to reports by [24]. For integration, first percept durations were found to be longer in the mean than subsequent percept durations (with statistical significance near equidominance; *p*-value of 0.0003 at *DF* =3 and 0.0184 at *DF* =5, right-sided Wilcoxon rank-sum test at significance level 5%). Mean durations of first *I*-percept were 10.9, 5.3 and 3.1 s, decreasing with *DF* = 3, 5, 7 (*p*-value of 0.0002 when comparing *DF* =3, 5 and 0.0014 for *DF* =5, 7). Mean durations of first *S*-percept were 3.5, 6.6, 8.1 s (comparisons did not produce statistically significant differences, possibly due to fewer instances of first *S* percepts). For subsequent percepts the means were the following: 5.4, 3.4, 3.1 s for *I* and 4.9, 5.2, 5.6 s for *S* at *DF* = 3, 5, 7, showing a decreasing trend for integration between *DF* =3 and *DF* = 5 or 7.

### B. Auditory streaming as an evidence accumulation process

Herein we propose an evidence accumulation model that accounts for the observed dynamical features of buildup and alternations: gamma-like distributions for first and subsequent durations, *DF* -dependent mean durations, and psychometric buildup functions. Data-based [1] estimates of spike counts of neurons in the primary auditory cortex (area A1) are sampled by a population of units and their summed responses lead to a population vote and to an increment of evidence “for” and “against” the current percept. When enough evidence has built up against the current percept, there is a switch to the opposite percept. Current increments can be positive or negative but only when the accumulated evidence is adequate, does a switch occur.

#### B.1. A basic state-dependent model for evidence accumulation

Our EVA model describes activity that accumulates and saturates at a target-level, *T*, just-subthreshold. The activity *X*_*n*_ is updated at the *n*th triplet according to:

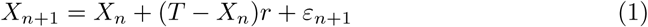

where *T* < 1 (assuming a unitary threshold) and where *ε*_*n*+1_ ∼ 𝒩 (0, *σ*^2^) are independent random variables (Gaussian noise of zero mean and standard deviation *σ*).

The activity increments are state dependent and proportional to the difference *T* − *X*, with constant rate *r*. Accordingly, the activity *X* drifts towards *T* stochastically if 0 < *r* < 1. Accumulation slows with *X*_*n*_ near *T* and the activity can cross the threshold only due to noise. At each threshold-crossing *X*_*n*_ is reset to a value *X*_*R*_ taken as the initial condition for the subsequent dynamics. The time *D* between successive threshold crossings represents a percept duration; it equals *N*_*D*_, the number of triplets between threshold crossings multiplied by the onset time from one *ABA*_ to the next (500 ms).

Phenomenologically, Eq (1) accounts for the features of the behavioral data described in Section A: the observed *DF* -dependent mean durations, the gamma-like shape of the distributions for first and subsequent durations, and the time course of the psychometric buildup. As an example, consider *DF* =5 and take *r*=0.6, which is the asymptotic, approximate value of the behavioral buildup, near equidominance (Fig 1C; red curve). With initial and resetting conditions *X*_0_ = 0.7 and *X*_*R*_ = 0.6, and with parameter settings *T* = 0.9, *σ* = 0.085, we simulated Eq (1) 675 times. The computed distribution of first *integration* normalized durations and the corresponding trial-averaged buildup function (Fig 2; right panels) are in agreement with Fig 1C-D.

**Fig 2.**
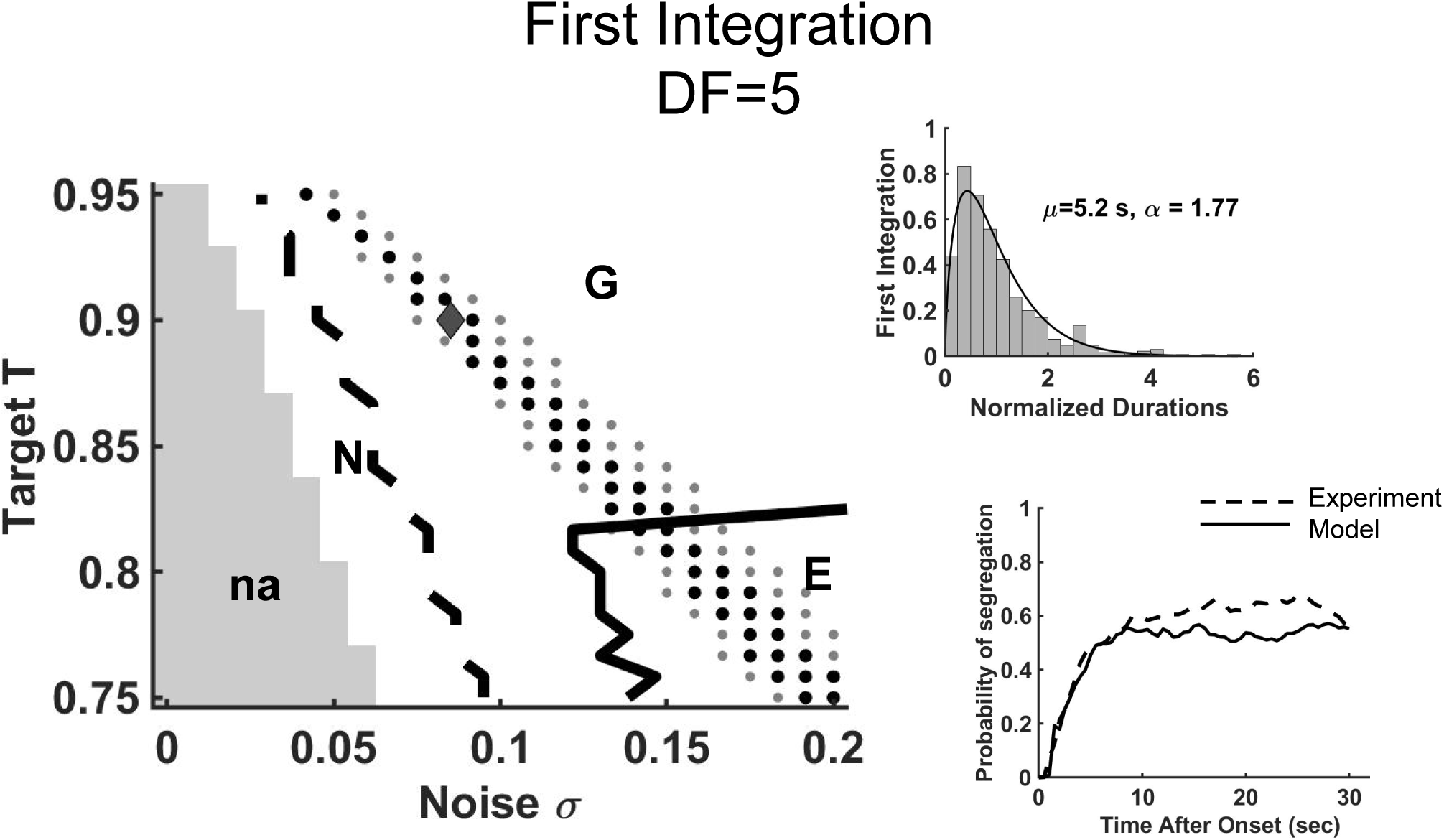
A basic state-dependent model for evidence accumulation yields percept durations that are gamma-like distributed and with mean values similar to those observed in behavioral data. To demonstrate the robustness of the model results and dependence on parameter values we simulated Eq (1) with various values for target *T* and noise level *σ*. Shown for *DF* =5 with *r* = 0.6: (Left) Two-parameter response diagram of the first *I*-percept with respect to *T* and *σ*. There is no switch for very small noise levels (na; gray area). Threshold-crossing activity appears with increased noise and leads to percept durations that are distributed according to normal distributions (region N), gamma-like distributions (region G between the black dashed and black solid curves), or exponential distributions (region E). Parameter values that lie on the sheets of black and gray dots yield numerically generated first integration mean durations within one and two standard deviation(s) of the experimental mean. (Right) Insets are shown for *T* = 0.9, *σ* = 0.085 (black diamond in the diagram): computed distribution of first integration normalized durations and the early phase of numerical buildup obtained during one simulation run of Eq (1) are in agreement with behavioral data. For simplicity, the drift rate *r* was kept constant to 0.6 between all threshold crossings.

To demonstrate the robustness of the model results and dependence on parameter values, we simulated Eq (1) with various values for target *T* and noise level *σ*. The simulations took into account the latency period during each trial, and the proportion of first percepts reported as integration and segregation, during the behavioral experiment at *DF* =5 (as in *Switches and resetting conditions*, in Methods). In this way we could assign a “percept”-type identity label to each event between consecutive resets. For any fixed *T* we found no threshold-crossings when *σ* was small. Alternations between “percepts” occurred only as *σ* increased, with dominant durations distributed as follows (e.g. for first *I*-percept; see Fig 2): normal distributions (region labeled N), gamma-like distributions with shape close to that found experimentally (region G) and exponential distributions (region E), respectively. The values of *σ* and *T* for which numerically generated mean percept durations were within one and two standard deviation(s) of the experimental mean were also calculated (Fig 2; sheets of black dots and gray dots). There is, therefore, a region in the parameter space where critical statistical properties of the behavioral data can be reproduced.

#### B.2. Linking neural data with behavioral data in the EVA framework

As in Micheyl et al. [1] we seek to relate perceptual buildup and bistability reported by human subjects during triplet-streaming to animal neural data. Spiking activity evoked by the *B*-tone was recorded from tone-*A*-selective neurons in macaque primary auditory cortex, A1 [1]. At any triplet position in the *ABA*_ sequence the mean spike counts, *m*_*DF*_, decreased with increased *DF*. The time course of *m*_*DF*_ exhibited fast adaptation and stabilized by the third triplet [1, Fig.3].

For modeling we assume that individual spike counts are Poisson distributed with means *m*_*DF*_, and that *m*_3_ > *m*_5_ > *m*_7_ for *DF* =3, 5, 7. The responses of A1 neurons will be processed by downstream neurons whose responses at each triplet then feed into the EVA accumulator. Suppose that *N*_*in*_ A1-neurons activate a neuronal unit downstream if the mean input exceeds a threshold *C*_*th*_

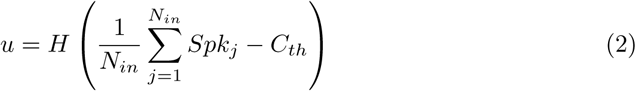

where *H*(·) is a Heaviside function. Such a sampler neuron is binary, taking a value *u* of 0 or 1 with probabilities *p* and 1 − *p*, for triplets after a brief transient phase of adaptation in A1. In line with the population separation hypothesis [7], when spike counts are large (so *u*=1) we tag the sampler as evidence for integration; likewise when spike counts are small (*u*=0), we tag the sampler as evidence against integration. As *N*_*in*_ increases, the averaged spike count variability around the mean *m*_*DF*_ decreases inversely with 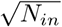 (Fig 3). As such, probabilities *p*_*DF*_ obtained from *m*_*DF*_ at different *DF* =3, 5, 7 vary from values being graded (when *N*_*in*_ is small) to values spread apart (when *N*_*in*_ is large), approaching extreme values of zero or one (when *N*_*in*_ is very large); Fig 3. A suitable variability in the A1-neuronal population can be chosen (more about this later in Sections B.4 and C.1) to ensure that probabilities *p*_*DF*_ at *DF* =3, 5, 7 achieve the graded asymptotic values of the behavioral buildup functions (e.g. 0.45, 0.6, and 0.65 as in Fig 1C). This is an important feature of the EVA model; indeed, without adequate variability in the readout of A1 responses we cannot account for graded BUF levels.

**Fig 3.**
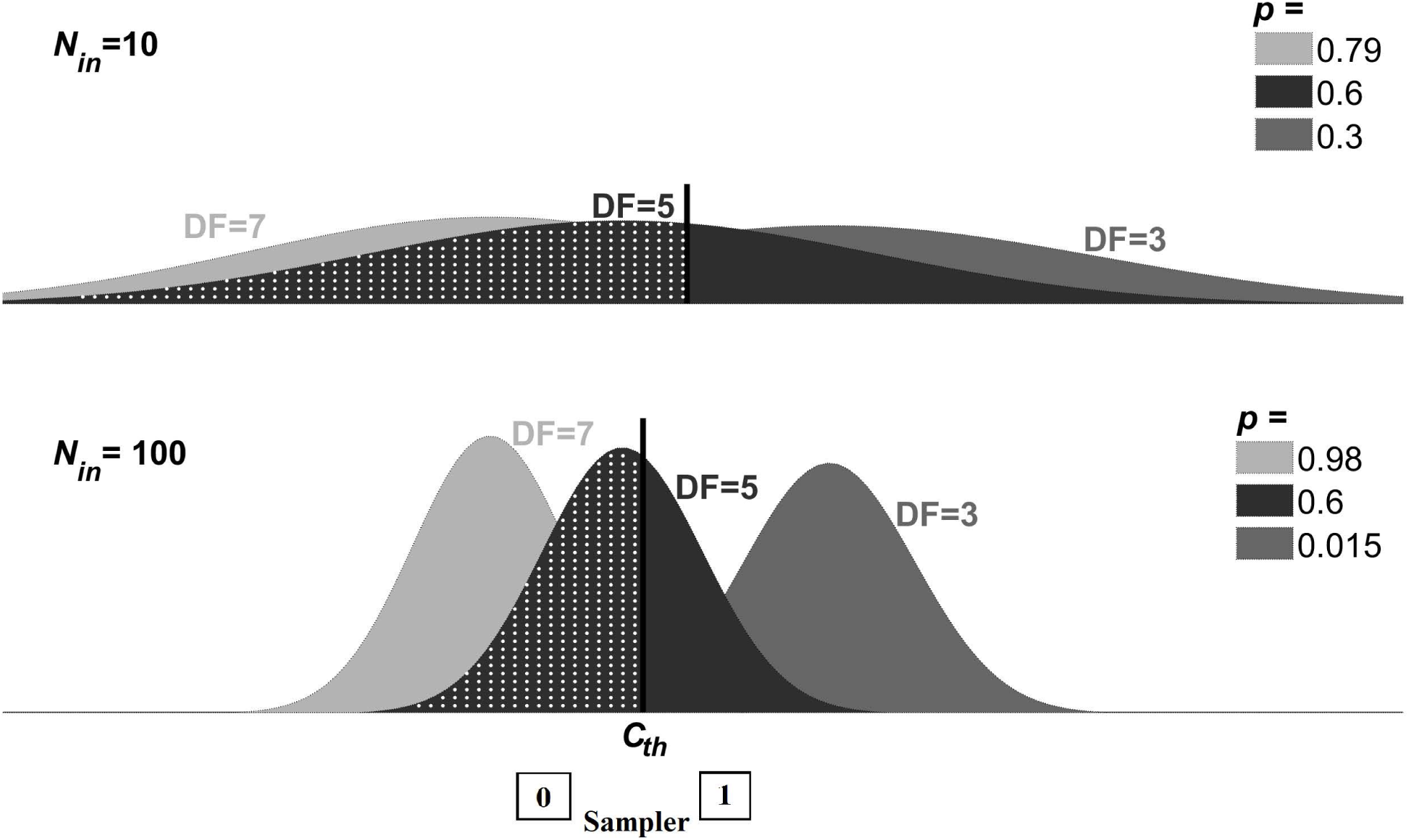
Linking neural data with behavioral data in the EVA framework. Individual spike counts of A1 neurons are assumed to be Poisson with means *m*_*DF*_ such that *m*_3_ > *m*_5_ > *m*_7_ for *DF* =3, 5, 7. Averaged spike counts over *N*_*in*_ A1-neurons,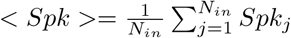, are normal-like distributed with means *m*_*DF*_ and standard deviation decreasing inversely with 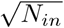; shown in gray (*DF* =3), black (*DF* =5) and light-gray (*DF* =7) for *N*_*in*_=10 (upper panel) and *N*_*in*_=100 (lower panel). At each triplet, < *Spk* > activates a sampler unit downstream if it exceeds a threshold *C*_*th*_ (solid black, vertical line). The area under the probability distribution to the left of *C*_*th*_ (white-dots pattern; *DF* =5) determines the probability *p* of the sampler neuron to be inactive (0); the complementary probability 1 − *p* is for the sampler to be active (1). For each *N*_*in*_, the threshold *C*_*th*_ was chosen such that *p* = 0.6 at *DF* =5, which is the asymptotic, approximate value of the corresponding behavioral buildup near equidominance (see Fig 1C; red curve). Probabilities *p* obtained at different *DF* vary from values being graded when *N*_*in*_ is small (*N*_*in*_=10), to values spread apart approaching zero or one when *N*_*in*_ is large (*N*_*in*_=100). A suitable variability in the A1-neuronal population is key if aiming to account for graded BUF levels observed in behavioral data (Fig 1C).

A question remains: How can we link the probability of a sampler unit becoming active stimulated by A1-neurons to the neuronal drive of the accumulator, *r* in Eq (1)? We resolve this problem by including an entire layer of binary units *u* as above, say *N*_*sl*_ total, and use the percentages *p*_*I*_, *p*_*S*_, as cluster sizes, of active and inactive samplers as input-drive to two accumulators: for and against integration, respectively (Fig 4A). In particular, the output *p*_*S*_ of the sampler layer (not binary anymore) is a stochastic process with mean *p* and variance *p*(1 − *p*)*/N*_*sl*_ (for justification, see *Statistical properties of SL-activation*, in Methods). Noteworthy, under this construction, the output *p*_*S*_ of the sampler layer (the input to the accumulator “against integration”) takes indeed values very close to *r*, defined as *p*, in Eq (1) if *N*_*sl*_ is large enough.

**Fig 4.**
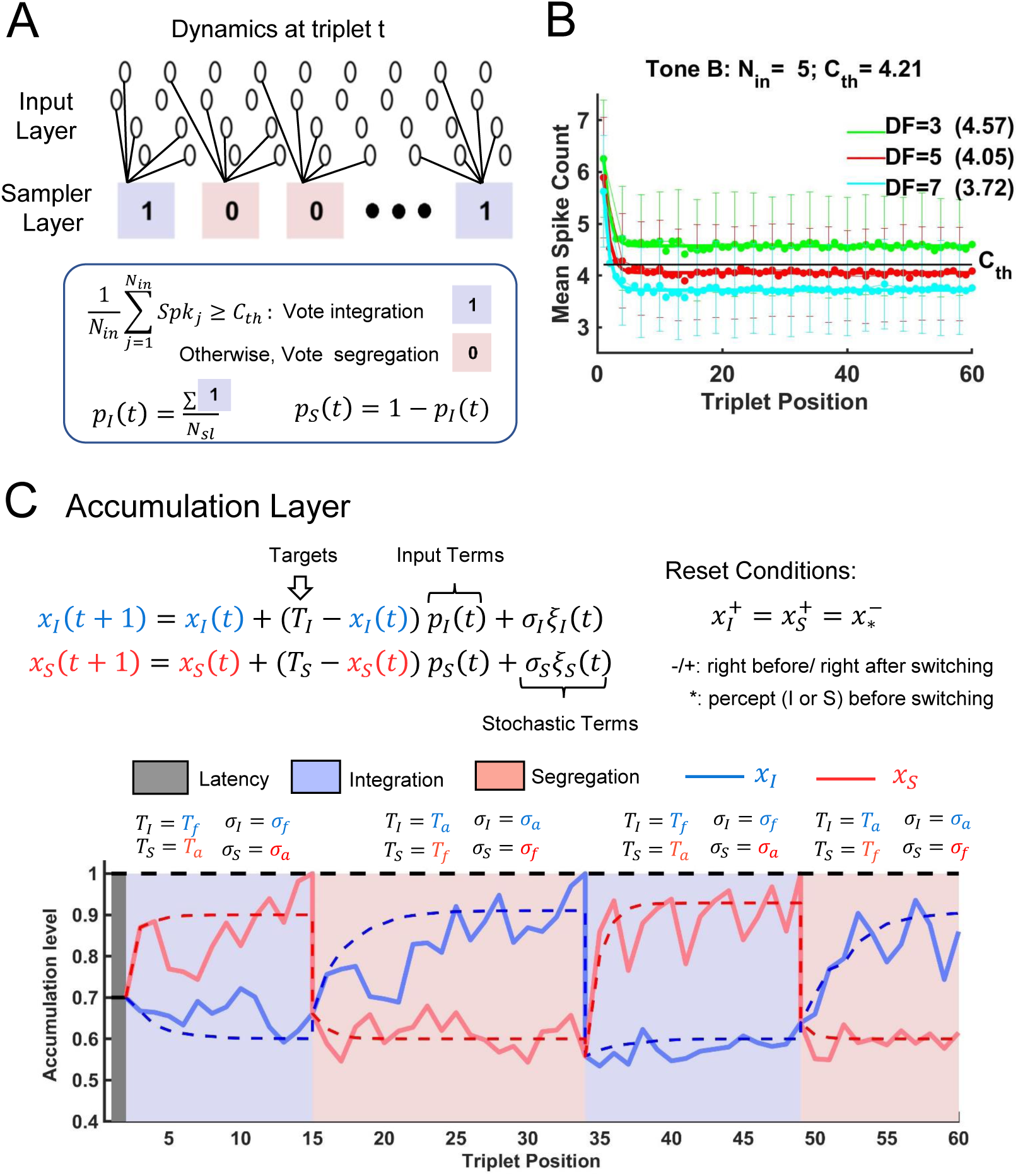
Accumulation model as feed-forward auditory network of 3 layers. A: State of neurons at triplet *t* in the input layer and sampler layer of the evidence accumulation model. Input layer comprises A1 units with (triplet- and *DF* -dependent) mean spike counts presented in panel B. Sampler layer has *N*_*sl*_=20 binary neuronal units, either in state 1 (blue; favoring *I* percept) or state 0 (red; favoring *S* percept). Each unit samples a small number of input units (*N*_*in*_=5) and the averaged spike count across the units is compared to *C*_*th*_ (see panel B) to determine the unit’s appropriate perceptual state. B: Mean spike counts (scatter plot) for tone *B* of tone-*A*-selective neurons, and exponential fit (solid) of mean spike counts. These values are interpolated for our specific *DF* =3,5,7 using data from cortical area A1 of awake macaque extracted from [1]. A Poisson spike count is generated using the mean value at each triplet. Asymptotic values of mean spike count (printed in parenthesis next to corresponding *DF* values) are used to generate spike counts after the 20-th triplet. Poisson spike counts are averaged across sets of *N*_*in*_=5 neuronal units, and the resulting values are subject to a binary neural threshold *C*_*th*_ (black horizontal line). The error bars indicate the standard errors of the mean spike counts. C: Accumulation layer has 2 accumulators drifting over successive triplets towards their own target values *T*_*a*_ and *T*_*f*_ where *T*_*a*_ > *T*_*f*_. Their activities are governed by input factors from the sampler layer and stochastic factors. The noise level depends on the target (*σ*_*a*_ > *σ*_*f*_). During a cycle, the suppressed unit accumulates evidence against the current percept. A switch to the other percept occurs when the accumulator of the suppressed unit reaches the switching threshold of 1. A new cycle starts, with accumulators reset to appropriate values, and targets values switched to corresponding perceptual states. Shown for *DF* =5. For other *DF* values, see S2 Fig. For the complete list of parameter values, see Methods – section *Parameter values used in model simulations*.

#### B.3. The EVA model

Our proposed EVA model is structured as a three-layer network (Fig 4; see details in *Methods*). It takes Poisson spike counts from tone-*A*-selective A1-neurons (the Input Layer, IL) [1, 7, 8] and passes them through binary units in the Sampler Layer, SL (Fig 4A). Only spike counts recorded during tone *B* are included (Fig 4B). Each SL-unit compares the averaged spike count across a small number *N*_*in*_ of input units to a fixed threshold *C*_*th*_ and places the outcome into either state 0 (for *S*) or 1 (for *I*). High activation in IL (above *C*_*th*_) is assumed to support percept *I* while low activation facilitates percept *S* (Fig 4A-B). The proportions *p*_*I*_ (*t*), *p*_*S*_ (*t*) (*p*_*S*_ = 1 − *p*_*I*_) of SL-units in states 1 and 0, together with stochastic noise terms *ξ*_*I*_ (*t*), *ξ*_*S*_ (*t*), modulate the activity of the Accumulation Layer, ACC (Fig 4C). Two accumulators representing evidence for the percepts drift towards two targets. Their activities *x*_*I*_, *x*_*S*_ are updated at discrete time steps determined by the position *t* of each triplet in the *ABA*_ sequence. One unit accumulates evidence for the current percept (e.g. *x*_*I*_ during integration) in the presence of additive “neural” noise defined by a Gaussian process of strength *σ*_*I*_ =*σ*_*f*_, and approaches target *T*_*I*_ =*T*_*f*_. The other accumulator works against the current percept (*x*_*S*_ during integration). It experiences stronger noise level *σ*_*a*_, and approaches another target, *T*_*a*_. Differential noise levels enable the accumulator “against” to be the first to reach the threshold and initiate the switch; meanwhile, the accumulator “for” remains confined to a neighborhood of its target. In the deterministic (noise free) case, alternations between percepts are not possible given that both *T*_*a*_ and *T*_*f*_ are subthreshold targets (*T*_*f*_ < *T*_*a*_ < 1), a distinctive feature of our accumulation model. Instead, the ACC system is bistable with accumulators *x*_*I*_, *x*_*S*_ reaching either steady state (*T*_*a*_, *T*_*f*_) or (*T*_*f*_, *T*_*a*_) depending on the initial conditions (Fig 4C, dotted lines in blue and red). In the presence of noise, however, the accumulator against the current percept reaches the decision threshold; a switch to the other percept occurs, the accumulators are reset, the targets are swapped (*T*_*S*_ =*T*_*f*_, *σ*_*S*_ =*σ*_*f*_ and *T*_*I*_ =*T*_*a*_, *σ*_*I*_ =*σ*_*a*_), then another accumulation cycle begins (Fig 4C, traces for *x*_*I*_, solid blue, and *x*_*S*_, solid red; the percept’s type is identified by the background color, blue for *I*, red for *S*. See also S2 Fig). It is essential that the accumulators are subjected to noise in order for the distribution of threshold crossing events idealizing the percept durations to exist.

#### B.4. EVA model captures *DF* -dependence of mean durations

Numerical simulations of the EVA model followed the experimental setup with *N*_*tr*_=675 repetitions (trials) per *DF*. The model-generated mean durations were computed separately for each percept type (*I, S*; first, subsequent) and *DF*. They approximated well their counterparts from behavioral data (Figs 5B and 6B). They also captured two important *DF* -related trends reported by other studies. First, near equidominance (*DF* =3, 5) mean durations of the first *I* percept were found to be longer than those of subsequent *I* percepts [2, 6]. Secondly, mean durations for *I* and *S* showed a “cross-diagram” like behavior [6, Fig.9B] with equidominance near *DF* =5. Mean durations for *I* were greater than mean durations for *S* when *DF* low (*DF* <5), and smaller than mean durations for *S* when *DF* large (*DF* >5), results similar to [6]. The model was robust to noise as demonstrated by 100 Monte Carlo runs of each *DF* simulation that yielded consistent results in terms of average values and 95% CI (Figs 5B and 6B; error bars).

**Fig 5.**
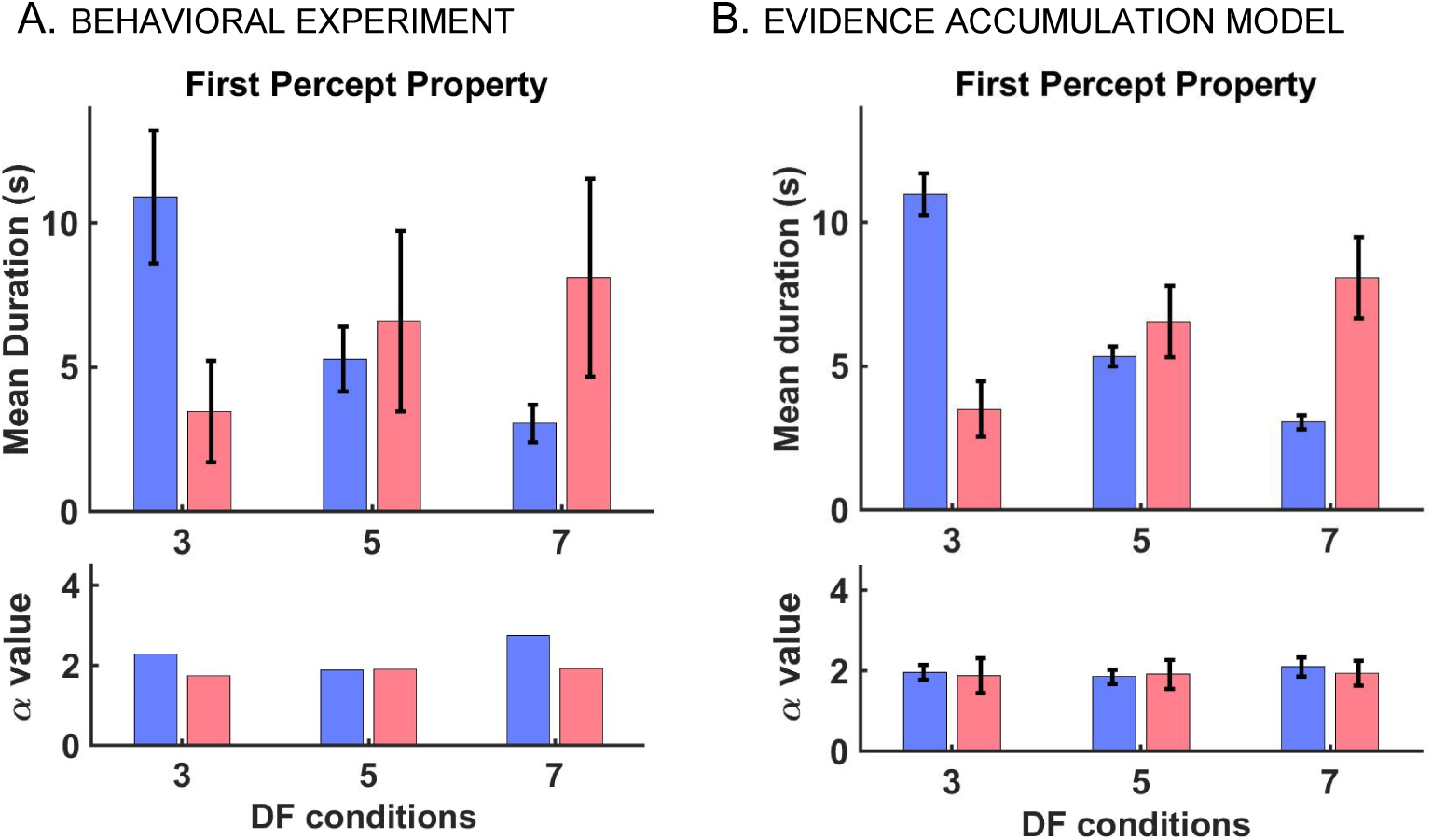
EVA model yields realistic first percept durations. A: Mean percept durations (top) and fitted *α* value (bottom) from gamma distribution of first *I* (blue) and first *S* (red) percepts from behavioral experiment for *DF* =3,5,7. The error bars indicate 95% CI around the mean and are obtained from statistical bootstrapping (see *Methods*). The mean durations of *I* decrease with *DF* while those of *S* increase with *DF*. The shape parameters *α* from gamma-fit using MLE for *DF* =3,5,7 are also presented here. There is no observed trend for *α* values. B: Mean percept durations (top) and fitted shape parameter *α* (bottom) from gamma distribution of first *I* (blue) and first *S* (red) states from EVA model. The error bars are 95% CI obtained from 100 Monte Carlo runs to show the robustness of the model. The mean values of duration follow the similar trend as those from experiment. Also, the shape parameters show a close resemblance to those from the experiment. Related results are included in S5 Fig.

**Fig 6.**
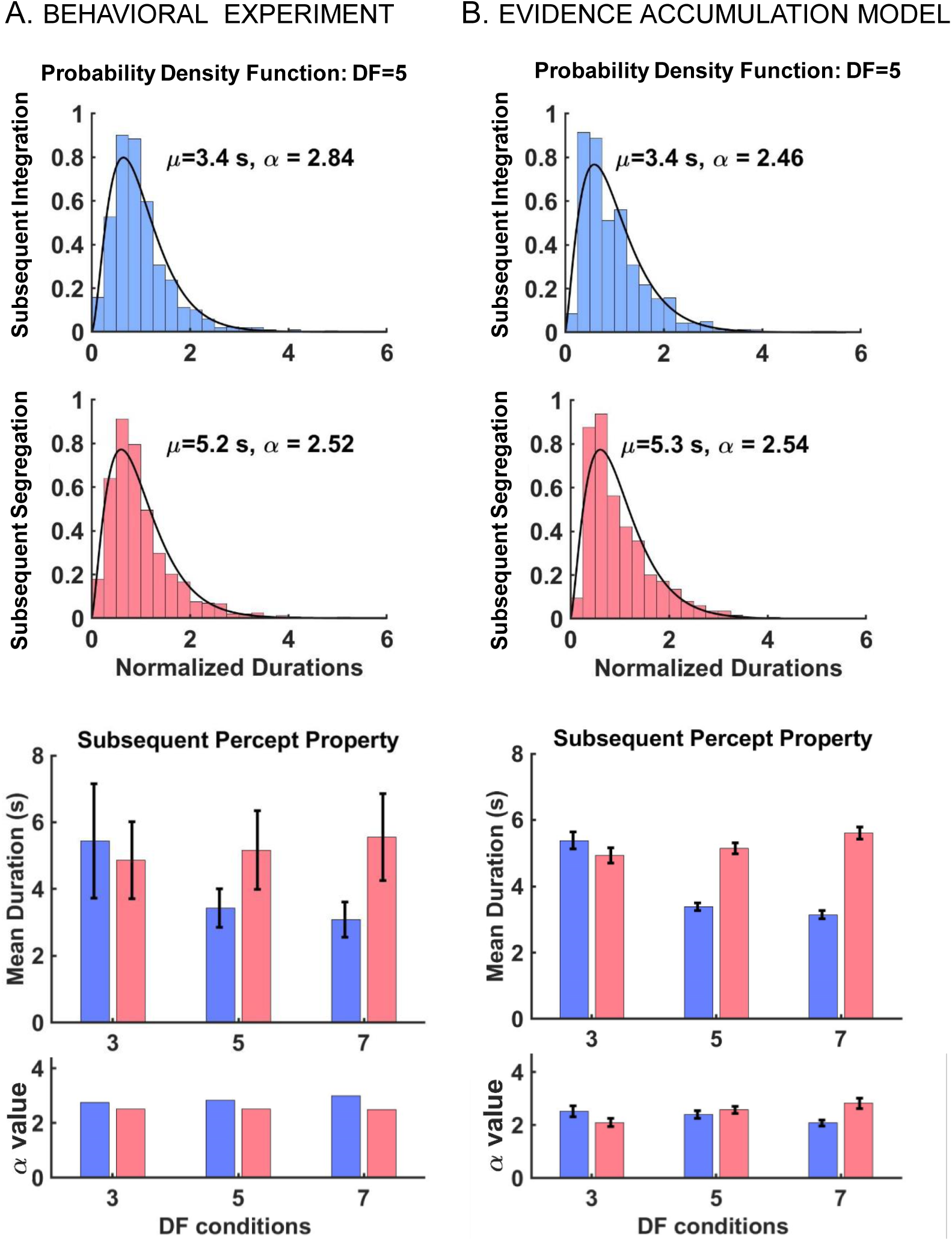
EVA model yields realistic subsequent percept durations and distributions. A: Distribution of normalized subsequent percept durations (top) and other properties (bottom) from behavioral experiment. (Top) Likelihood ratio test confirms that both subsequent *I* (blue; *N* =1642, *p*=0.49) and *S* (red; *N* =1785, *p*=0.49) percepts follow a gamma distribution, shown here for *DF* =5. The shape parameters *α* computed using MLE, and the mean durations *µ* are shown in the graphic. (Bottom) Mean subsequent durations for *I* (blue) and *S* (red) for *DF* =3,5,7. The error bars indicate 95% CI around the mean, computed using statistical bootstrapping (see *Methods*). Similar to the first percept, mean durations of subsequent *I* and *S* show a “cross-diagram” like behavior [6, Fig.9B] with equidominance near *DF* =5; the ratio between mean durations for *I* and *S* percepts changes from larger than 1 to smaller than 1 when crossing *DF* =5, near equidominance. The shape parameters *α* from MLE for *DF* =3,5,7 are also presented, and no trend for *α* values is found. B: Distribution of normalized subsequent percept durations (top) and properties (bottom) from EVA model. (Top) Normalized subsequent percepts are gamma distributed for *DF* =5 with mean durations *µ* and shape parameters *α* shown in the figure; similar results are obtained for other *DF* values, see S1 Fig. (Bottom) Mean percept durations and fitted shape parameters *α* for *DF* =3,5,7 from EVA model. Mean subsequent durations follow the same trend and the shape parameters have similar values as compared to those from the experiment. The error bars are 95% CI obtained from 100 Monte Carlo runs of the model. The result shows the robustness and consistency of the model. Related results are shown in S5 Fig.

**Fig 7.**
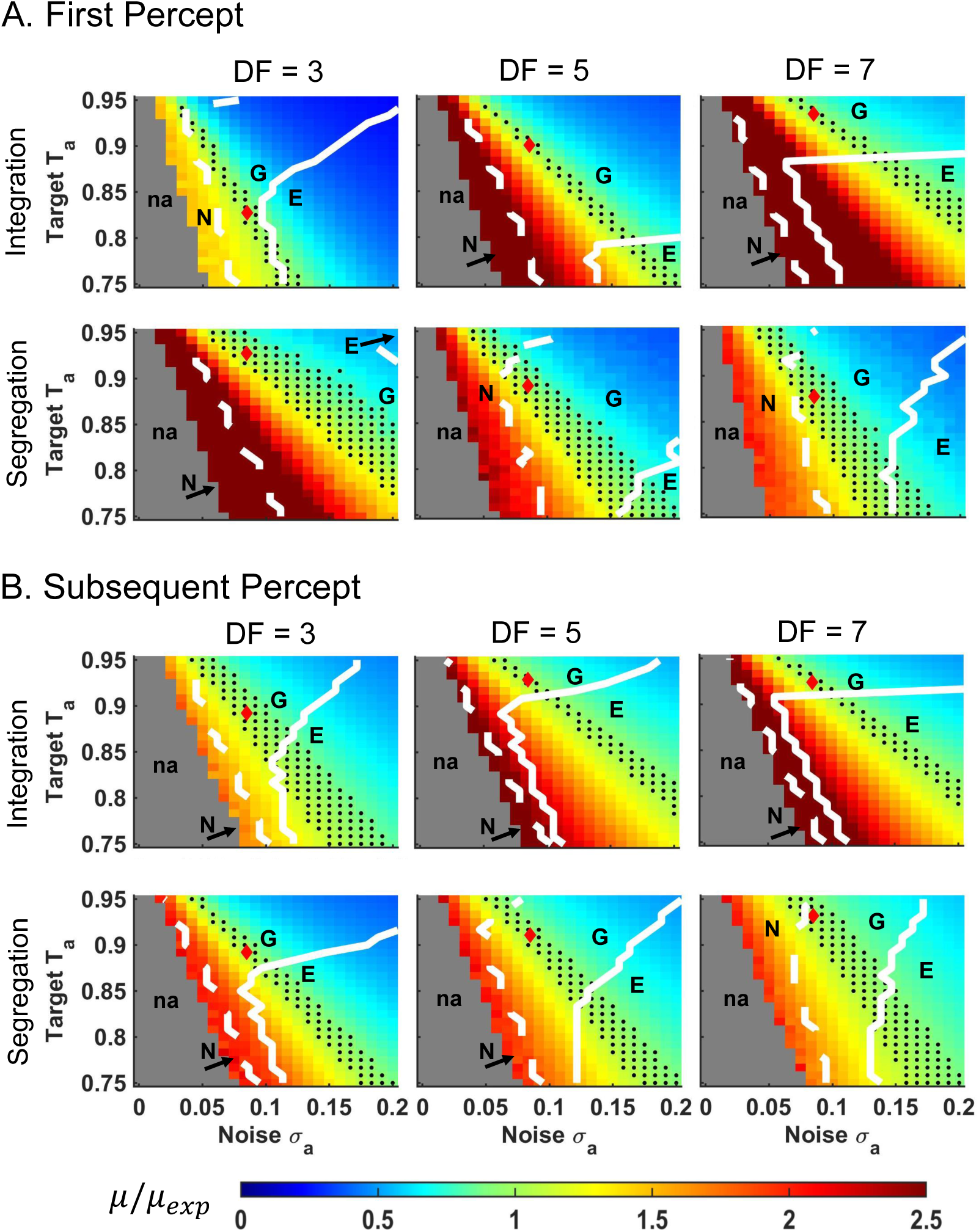
Dependence of mean percept durations and shape of distributions on target *T*_*a*_ and stochastic term *σ*_*a*_, in the EVA model. Diagrams show the difference between the mean duration *µ* derived from numerical simulations of the model and mean *µ*_*exp*_ from the behavioral data, represented as ratio *µ/µ*_*exp*_; see the color scheme. Results are shown for A: first and B: subsequent integration and segregation percepts at conditions *DF* = 3,5,7. Given a fixed value for *T*_*a*_, the dynamics changes from no alternations between percepts at small *σ*_*a*_ (na; in gray); to alternations of mean durations much longer than the experimental mean (region in warm colors); to mean durations that approximate well the corresponding experimental values (within one standard error to *µ*_*exp*_; in green; black dots depict a discrete selection of values in the green region); then, to mean durations much shorter than *µ*_*exp*_ (region in cool colors). In each diagram, *σ*_*a*_, *T*_*a*_ that were used to generate model-based results are identified by a red diamond (see Methods, *σ*_*a*_ = 0.085, *T*_*a*_ varies). Besides mean durations, the shapes of the gamma-like distributions that fit normalized percept durations depend on *T*_*a*_ and *σ*_*a*_ as well (*α* is the shape-parameter in the gamma-fit; see Eq (3) in Methods). There are three main regions that characterize *α* and they are delineated by the dashed-white and solid-white curves. Low-level of noise *σ*_*a*_ yields normal distributions (region N, to the left of dashed-white line; *α* ≫ 3) while high-level of noise yields exponential distributions (region E, to the right of solid-white line; *α* near 1). For intermediate level of noise, the distributions are gamma-like with shape close to that found experimentally (region G, between white contours; *α* ≈ 2 for first percepts and *α* ≈ 2.6 for subsequent percepts; *α* differs from *α*_*exp*_ by relative error up to 20% except for integration at *DF* =7 where it is up to 30%). The parameter range where both model-generated mean duration and shape of distribution are good approximations of their corresponding experimental observations is found at the intersection of region G with the sheet of black dots. Related results are shown in S3 Fig.

**Fig 8.**
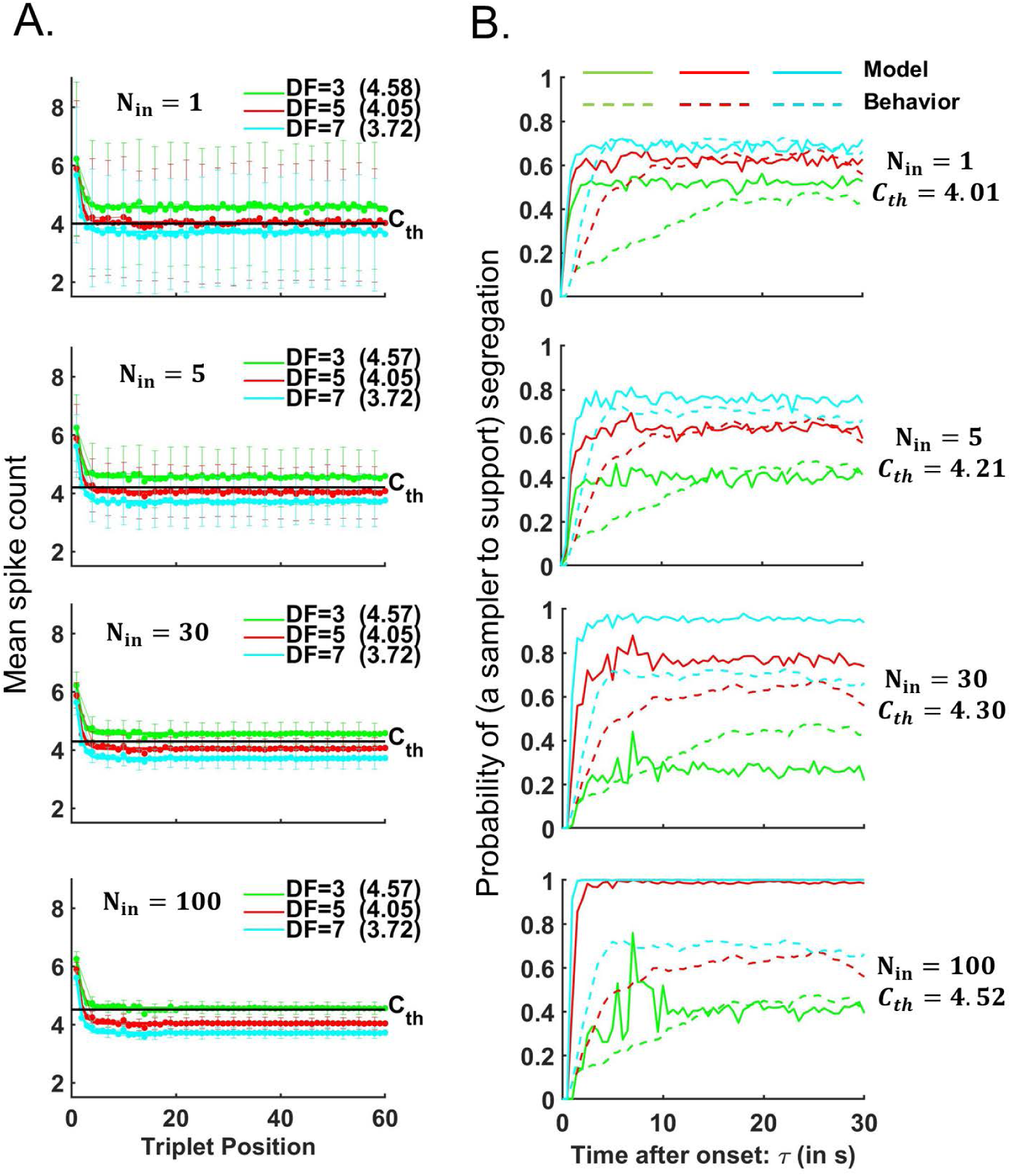
Parameter fitting for input and sampler layers in the EVA model. The signal detection algorithm for constructing a neurometric function (the probability of a sampler to support the segregation percept) utilizes spike count time courses as shown in panel A (data extracted from [1] and interpolated for the cases *DF* =3, 5, 7); see below for more detail. The behavioral buildup functions (dashed, in panel B) occupy intermediate ranges of probability of *S*, and show slow initial rise for *DF* =3, 5. The simulated functions (solid, in panel B) do not capture the slow-rising phase of behavior buildup, and the spread between the neurometric curves increases unacceptably at larger *N*_*in*_. For an optimal choice of parameters *N*_*in*_, *C*_*th*_, the algorithm yields well-fit asymptotic values of behavioral data. A: Mean spike counts *m*_*t*;*DF*_ are interpolated at *DF* =3, 5, 7 st from data in [1], and then extended for triplets *t ≤* 60; see Methods. (Note: In [1, Fig.3] mean spike count data were shown for *A*-tone selective neurons in A1 during triplet tones at *DF* =1, 3, 6, 9. They decreased exponentially and stabilized within a few seconds. Mean spike counts changed with *DF* only during *B*-tone.) Herein, spike counts during *B*-tone are generated using Poisson processes of means *m*_*t*;*DF*_ (*DF* =3, 5, 7) and then averaged over *N*_*in*_ neuronal units of the input layer IL (e.g. *N*_*in*_= 1, 5, 30, 100). The average values of the mean spike counts and the standard error to the mean (SEM) are computed over 675 trials. Averages, including asymptotic values (written in parenthesis, at each *DF*), do not change with *N*_*in*_ but SEM decreases with a factor of 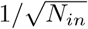. B: The signal detection algorithm [1] generates neurometric functions using numerical data from IL-pools of *N*_*in*_ neuronal units; parameter *C*_*th*_ is chosen to yield the least-squares error of the experimental buildups and the computer-simulated neurometric functions for *DF* =3,5,7. If *N*_*in*_ is small the neurometric curves tend to bunch together due to overlapping and large SEM regions across conditions. As *N*_*in*_ gets bigger, the neurometric curves are pushed apart. The best approximation to the set of psychophysical buildups is obtained for *N*_*in*_ = 5, *C*_*th*_ = 4.21.

**Fig 9.**
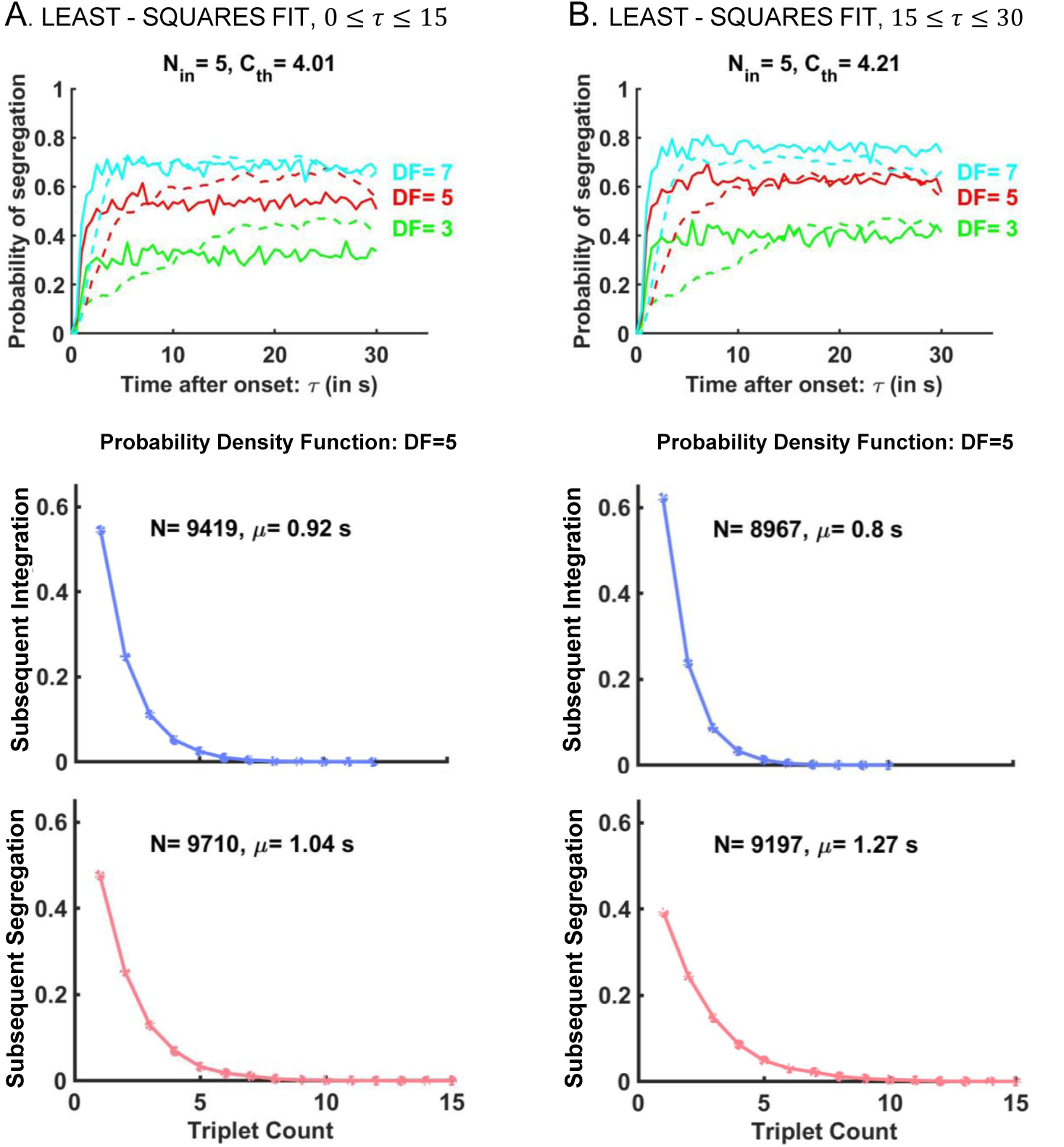
Signal detection algorithm adapted from [1] yields exponential distributions and unrealistic mean durations of percepts. (Top) Binary threshold *C*_*th*_ is chosen to yield the least-squares error between neurometric buildups (solid) and behavioral buildups (dashed) at *DF* =3,5,7. Poisson spike counts are averaged across a sample of *N*_*in*_=5 neuronal units and compared to *C*_*th*_ to classify a triplet either as *I* or *S*. Trial-averaging the *S*-tagged responses produces the neurometric functions. The threshold value is determined by least-squares fit for A: the first 15 seconds of the stimulus to match the transients, or for B: the last 15 seconds of the stimulus to match the asymptotic level of the behavioral buildup; See also Fig 8. (Bottom) Trial-by-trial applications of the signal detection algorithm from [1] with A: *C*_*th*_=4.01 and B: *C*_*th*_=4.21 yield exponentially distributed subsequent percept durations for *I* (blue) and *S* (red). Their mean values *µ* are significantly smaller than those reported in the experiment. Note: Same parameter values *N*_*in*_=5, *C*_*th*_=4.21 were used in the EVA model for activation of the sampler layer SL (Fig 4A-B) and obtain gamma-like distributions of percepts (Figs 1D and 6B).

In EVA model, the switch to a new accumulation cycle occurred when the ACC-unit that accumulated evidence against the current percept reached the decision threshold. Target-against *T*_*a*_, noise level *σ*_*a*_, and increment rate *p*_*S*_ (*t*) determined the trajectory of the suppressed unit *x*_*S*_ and the length of the corresponding dominant percept *I*. Similarly, *T*_*a*_, *σ*_*a*_ and *p*_*I*_ (*t*) determined the duration of percept *S*. We studied the effect of *T*_*a*_ and *σ*_*a*_ on the model-generated mean durations at each *DF* by varying their values while keeping all other parameters fixed (see *Parameter values used in model simulations*, in Methods). At a given *T*_*a*_, EVA model exhibited no alternations if *σ*_*a*_ was small (Fig 7; region in gray). For moderate *σ*_*a*_ values, perceptual switches occurred but yielded percepts of mean durations much longer than those found experimentally (in warm colors); then, for larger *σ*_*a*_, simulated durations became comparable to (in green) and then much shorter than (in cool colors) the behavioral mean durations. Similar results were obtained for *σ*_*a*_ fixed when varying *T*_*a*_. As a general rule, the smaller the target-against, the stronger the noise level had to be in order for the accumulator’s trajectory to be pushed above the threshold and to generate acceptable statistical approximations of the data (Fig 7, in green; also black dots; within one standard error to the experimental mean, SEM).

The decrease in mean durations of *I* percepts with increasing *DF* (Figs 5B and 6B, blue) stemmed from the increase in probability of a sampler to support segregation (Fig 8B) which led to an increase of increment rate *p*_*S*_ of accumulator *x*_*S*_ (see *Statistical properties of SL-activation*, in Methods). The increasing trend of mean durations of *S* percept, with *DF*, could also be associated with the decrease of increment rate *p*_*I*_ of accumulator *x*_*I*_. These *DF* -dependent properties of *p*_*S*_, *p*_*I*_, inherited from A1 spike counts (Fig 8A) enabled the EVA model to capture the correct qualitative trend of the experimental means across all percept-types and *DF* conditions. Suitable quantitative agreements were then obtained by fine-tuning the value of target-against *T*_*a*_ (Fig 7, red diamond; error between numerical and behavioral results was restricted to 0.1 SEM. See also S3 Fig).

#### B.5. EVA-modeled first and subsequent percept durations match observations

The model reproduces an important statistical feature of the behavioral data, the distributions of normalized durations for all first and subsequent *I, S* percepts at *DF* = 3, 5, 7. Histograms were drawn and fitted by gamma probability density functions of shape parameters *α* (see Eq (3) in Methods) whose values agreed with those from the behavioral experiment. The shape of distributions was tested and confirmed statistically by 100 Monte Carlo runs of the model for each *DF* condition separately (Figs 5B and 6B; error bars indicate 95% CI around *α*-mean values). Exemplar distributions for first and subsequent durations are shown in Figs 1D and 6B at *DF* =5. For other *DF* values, see S1 Fig.

Since alternations were caused in the model by the accumulator that gathered evidence against the current percept, the distribution of threshold crossing event times depended on *T*_*a*_ and *σ*_*a*_. In particular, for a given *T*_*a*_, EVA model generated percept durations that were normally distributed for *σ*_*a*_ small (Fig 7, region N; *α* was much bigger than 3) and exponentially distributed for *σ*_*a*_ large (Fig 7, region E; *α* was close to 1). For intermediate *σ*_*a*_, the distributions were gamma-like matching those fit to the observed data (Fig 7, region G; model-generated *α* values were similar to those determined from experiments, *α*_*exp*_, at relative error up to 20-30%). The closer *T*_*a*_ was to the decision threshold 1, the easier it was to find *σ*_*a*_ that yielded gamma-like distributions. With decreasing *T*_*a*_, the transition to a narrower region G was either sharp-edged (e.g. *DF* = 7, first *I*) or rather smooth (*DF* = 7, first *S*). Percept durations that approximated well both the distribution shape and the mean duration of the experimental data were obtained by using parameters from region G that overlapped with the black dotted sheet.

#### B.6. EVA model captures *DF* -dependence of stream segregation buildup

The model-generated buildup functions captured both the rising and the asymptotic phases of the behavioral buildup for each *DF* = 3, 5, 7 (Fig 1D). These trends were a consequence of already having simulated percept durations and percept means well-fit to behavioral data, in accordance with previous works describing the buildup of stream segregation as a byproduct of an alternating renewal process [17].

#### B.7. Computational advantages of the EVA model

The model is pseudo-neuromechanistic; it takes A1 responses as input, it allows for attractor-states, and it includes accumulators that are saturating akin to synaptic currents. The spike counts are in accordance with neurophysiological data from A1 [1] and provide input to the computation of perception dominance downstream, as in the conceptual population-separation model of [7] and in competition-based model of [6]. The model incorporates fast habituation (after one triplet or so, Fig 4B) as in [1, Fig.3] and it accounts for the decrease in response amplitudes and in spatial activity patterns evoked by tone *B* at tone *A* tonotopic locations observed as *DF* increases [1, 7]. Indeed, if most of tone-*A*-selective A1-neurons are active, the model predicts a large proportion of samplers in SL to be active (*p*_*I*_ large) and thus favors *I* percept. If the opposite happens and A1 is mostly inactive, a large proportion of samplers are inactive (*p*_*S*_ is large) and the model favors *S* percept. Activation in IL decreases with larger *DF* (fewer IL-units have mean spike counts above threshold *C*_*th*_) and so does *p*_*I*_ (*t*); Fig 4A-B, compare *DF* = 3, 5, 7; see also [1, Fig.3] and [7, Fig.11]. This affects the dynamics of the accumulators since *x*_*I*_ (*t*), *x*_*S*_ (*t*) gather evidence about percepts with increment rates proportional to *p*_*I*_ (*t*), *p*_*S*_ (*t*) respectively, while also being modulated by a certain level of noise (Fig 4C, Eqs).

The accumulators resemble discrete time versions of the leaky integrate-and-fire neuron model with conductance-based synaptic input [25], *dV* = (*V*_*R*_ − *V*)*Ddt* + *noise*. The voltage-like variable (here, normalized, by the threshold value for switching) has a maximum amplitude of one. The reversal potential *V*_*R*_ is set to either target *T*_*a*_ or *T*_*f*_ depending on the type of the dominant percept and the type of the accumulator. The synaptic drive *D* consists of feedforward input from SL and is analogous to the reciprocal of the time constant. Finally, Gaussian white noise represents input from other brain sources or internal to ACC. Its strength *σ* needs to reach an appropriate level for the statistics of percept durations generated by the EVA model to match the behavioral data (see *Methods*).

Our EVA model is data-driven. Initial conditions are set using the latency periods and the proportion of first *S*-percepts from the experimental data. Poisson spike counts of IL neuronal units at each triplet *t* and semitone difference *DF* are generated using mean values *m*_*t*;*DF*_ derived from macaque A1 multi-unit spiking neural data [1] (Fig 8A). Parameters *N*_*in*_ and *C*_*th*_ are obtained by least-squares fit between the probability of a sampler to support an *S* percept and the behavioral buildup at all three *DF* values (Fig 8B; see also *Methods*). Neuronal granularity as a suitable substrate for perceptual representations [24] is implemented through SL. The number of samplers is chosen flexibly from a wide range of values (*N*_*sl*_ ≥ 1; here *N*_*sl*_=20). There are few other free parameters, *T*_*a*_, *σ*_*a*_, *T*_*f*_, *σ*_*f*_, *b* (baseline), but only the former two are major players in fitting the model to data (as shown in Sections B.4 and B.5).

### C. Signal detection algorithm yields fast buildup and unrealistic percept durations

#### C.1. Modeling the buildup with Micheyl’s model for auditory streaming

The signal detection algorithm of Micheyl et al. [1] (see also [26, 27]) has been extensively cited in the auditory streaming research [3–6, 12, 21] in regard to computing time-varying probabilities of stream segregation from neuronal responses in A1. The model was based on choosing a threshold number, *C*_*th*_, for mean spike count (first, trial-averaged; then averaged across sampled neurons) to classify each triplet as *I* or *S*; doing this for each *ABA*_ in the sound sequence generated a time course, a “neurometric” function. Briefly, for a given triplet position, a probability distribution was constructed for the *B*-tone responses measured at and convolved over A1 neurons whose best frequency was that of the *A*-tones. The area under the probability distribution to the right of *C*_*th*_ determined the probability that tone *B* was detected and, consequently, that the triplet belonged to *I* percept; the complementary probability was associated with *S* percept. The algorithm classified each triplet independently and assumed no memory among nearby triplets.

A simplified view of this procedure is to consider the distribution of trial-averaged counts for the neurons as straddling the mean for each triplet in the time course of an *ABA*_ sequence [1, Fig.3]. Conceptually, one chooses a level *C*_*th*_ that will cut across the distributions, say for *DF* =3, and correspond to low probability of *S* for early time and correspond to the asymptotic level (from behavior) for late time (e.g. *C*_*th*_, thin horizontal line, in S4 Fig). This classification could likely provide a decent fit for *DF* =3 but for *DF* =6 the spike counts will fall below the threshold, leading to an overestimate of probability of *S*. The remaining cases will yield extreme classifications: for *DF* =1, spike counts for each triplet in the sequence will be above *C*_*th*_ and probability of *S* will be estimated as near zero; for *DF* =9, all spike counts will be below *C*_*th*_ (except maybe for the initial triplet) and therefore probability of *S* will be near one for the time course. The spread of the behavioral time courses in [1, Fig.4], one lying intermediate for *DF* =3 and the others at very low or quite high levels, provided an opportunity for reasonable fitting with a single *C*_*th*_ level [1]. For details on numerical fitting with such signal detection algorithm, see S4 Fig.

In the case of our data, the conditions were *DF* = 3, 5, 7 and the behavioral buildup functions lay in more intermediate levels and clustered around the asymptotic value of 0.5 (Fig 1C; also Fig 8B, dotted lines). It thus became challenging to fit the buildup functions (especially the early, slower rising portions for multiple *DF* values, 3 and 5 semitones) using Micheyl’s model with a unique *C*_*th*_.

We attempted to meet this challenge by applying the signal detection algorithm to our behavioral data while using interpolated and Poisson distributed spike counts based on the neural data from [1]. The approach was equivalent to the computation of the probability of a sampler to support segregation from the EVA model, observing only the input layer, IL, and passing its output through one single sampler, *N*_*sl*_=1. While the mean spike counts over a pool of *N*_*in*_ A1-neurons did not change significantly with *N*_*in*_, the standard error to the mean did (Fig 8A). The decrease in the spike count error to the mean made the horizontal line *C*_*th*_ intersect fewer local distributions and biased the data-fitting towards the behavioral curve for a certain *DF*, at the expense of others. Choosing more A1 neuronal units (larger values of *N*_*in*_; Fig 8A) led to larger spread in the simulated neurometric functions and poorer fitting (Fig 8B; at *DF* = 5, 7 the neurometric functions (solid lines) plateaued at probability approximately 1 after triplet-sequence onset, as *N*_*in*_ increased; e.g. case *N*_*in*_=100). The best approximation of the asymptotic levels of the behavioral buildup for all *DF* conditions was found at a relatively low *N*_*in*_ however the rising transients of the neurometric functions were still much faster than in the experiment.

#### C.2. Modeling percept durations with Micheyl’s model

We extended the work from [1] by using the signal detection algorithm to generate “percepts” and characterize their distributions. For each *DF*, adjacent triplets of the same type (*I* or *S*) were grouped together to create percept phase durations and construct frequency graphs (Fig 9). Theoretical calculations showed that subsequent percept durations generated by Micheyl’s model were exponentially distributed, as opposed to gamma-like. During subsequent durations we could assume that the buildup functions of stream segregation had reached an asymptotic level *p* (Fig 9A-B, upper panel) and that the activity in the A1 pool was independent at each triplet. Then the probability that percept *S* consisted of *n*-triplets could be calculated as *Prob*(*D*_*S*_) = *p*^*n*^(1 − *p*) = (1 − *p*)*e*^*n* ln *p*^, depending exponentially on *n*. Likewise the probability that *I* consisted of *n*-triplets was *Prob*(*D*_*I*_) = *p*(1 − *p*)^*n*^ = *pe*^*n* ln(1−*p*)^. Similar results were obtained from numerical simulations of Micheyl’s model. The probability density functions were found to be discrete versions of exponential curves and the mean durations were small at about 1 s (Fig 9A-B, middle and lower panels), suggesting that the signal detection algorithm is not appropriate to describe perceptual alternations and percept durations, key aspects of bistable stream segregation. (Compare to Fig 1C; also S1 Fig and [2, 5, 6].)

## Discussion

We developed a new evidence accumulation model for auditory streaming of triplet sequences *ABA*_*ABA*_ *…* that takes as input neuronal responses of primary auditory cortex, A1 (macaque, [1]). Our neural-like model accounts for the (human) behavior we observed under three conditions (tone frequency difference, *DF*). During trials, subjects reported spontaneous alternations (bistability) between integration, *I*, and segregation, *S*. The first percept was usually *I*; the probability of *S* built up over time rising from near zero and plateaued within a few seconds to a level that increased with *DF*. In the model, switching between *I* and *S* occurred when noisy accumulation of evidence against the current percept exceeded threshold. Our simulations matched both buildup time-courses and percept-duration distributions.

Our model draws inspiration from the population separation hypothesis of [7] and focuses primarily on the *B*-tone responses of *A*-tone selective neurons. Micheyl et al [1] used similar principles to compute “neurometric” functions for segregation buildup. Their signal-detection model was applied to A1 and to sub-cortical neuronal spike count data to conclude that perceptual organization of auditory streams was present in early stages of the auditory pathway [3, 21]. It treated each triplet as independent of the previous ones, without an accumulation process from triplet to triplet. The only time dependent mechanism was adaptation of A1 neurons that was nearly complete after 2-3 triplets – too fast to account for buildup. Herein we show that Micheyl’s model behaves as if classification is like coin-tossing with possible bias. Simulated durations are therefore like run-lengths in coin-tossing, exponential-like and very brief, contrary to the observed data (gamma or lognormal-like).

Our approach underscores the essential significance of duration distributions as characterizations of streaming and switching, a constraint overlooked by previous analysis [1]. It emphasizes that neuronal-based modeling of behavioral data that goes beyond trial-averaged behavior may need to involve an evidence accumulation process in order to account for the statistics of single trials.

### Novel features

Our model is intuitively straightforward. It describes the accumulation of evidence, incremental from each triplet, for or against the current percept. The estimated A1-spike counts are passed through a sampler layer, SL, each of whose units sample a few A1-neurons. SL-units vote 1 or 0 if the summed spike counts for the current triplet are above or below threshold. The fraction *p*_*I*_ (*p*_*S*_) of sampler-votes 1 (0) represents the net output which favors integration (segregation), transmitted to the accumulators. After multiplicative weighting, *p*_*I*_, *p*_*S*_ are used together with additive noise to update the accumulators. Of significance, the weighting factor is state dependent, proportional to the difference, *T* −*x*, between the current accumulator value *x* and a target *T*. Accumulation slows when *x* is closer to *T* and, importantly, we can choose *T*<1 in which case our model mimics noise-driven attractor competition dynamics [23]. Further, if *T* is close to one (i.e. accumulation saturates below, near threshold), gamma-like distributed threshold-crossing times are more robustly obtained with modest noise levels [22, 28].

State-dependent dynamics of stochastic accumulators in the framework of bistable perception were highlighted in other previous works [24, 29]. Our approach implements several distinctive features: a link to spike count neural data, an intuitive equation for the accumulator (see Eq 1, basic model for behavior), and a theoretical framework that goes beyond equidominance by looking at graded responses across multiple stimuli conditions.

Our model is a hybrid: it incorporates some neuro-based phenomenology (A1 neuronal responses as input, saturating driving force, escape dynamics) but it is non-committal to specific neuronal mechanisms of inhibition and adaptation. Moreover, key parameters are not directly linked to neuromechanistic processes but rather determined by fitting model dynamics (simulated threshold-passage times) to observed duration distributions.

### Duration distributions underlie buildup

Buildup functions (BUFs) for behavioral data are based on trial-averaging of ongoing reporting of percepts; the buildup functions can be well-reproduced by an alternating renewal process applied to the percept duration distributions [17], in spite of disregarding the small inter-duration correlations. Our EVA model, using neural data as input, as well reflects a choice process, a neuronal computation, based on single-trials. From the EVA-simulated switch times we computed the “percept” duration distributions and generated BUFs that compared well with the behavioral data. The single-trial percept durations are the critical observations for a model to match in order to characterize streaming dynamics for stimuli with constant parameter values such as *DF*. We conclude that trial-averaging of the spike counts, especially from too early in the cortical pathway, and a triplet-based signal detection scheme [1], washes out the dynamical aspects of accumulating neuronal computations that underlie perceptual multi-stability. Model-based analyses of trial-averaged neuronal responses that show ramping behavior in decision-making tasks have recently come under scrutiny by consideration of single-trial data [30]. Arguments were made, admittedly still under debate [31, 32], that trial-averaged smooth time courses of evidence accumulation during decision-making might arise from temporally “discrete steps” rather than from continuous ramping dynamics. We suggest that care be exercised when making interpretations from trial-averaged neuronal responses, neuronal ramping or neuronal BUFs, to consider that such averaging may overlook the discrete event nature of perceptual switching and/or decision-making that involve evidence-accumulation/competition.

### Fitting of model to data

We assigned different values of noise and targets to “against” and “for” accumulators to ensure switching was caused by the against-unit crossing the decision threshold. Intuitively, as the accumulator-against saturates around target-against *T*_*a*_ (subthreshold), enough noise *σ*_*a*_ guarantees threshold-crossing. The closer *T*_*a*_ is to one, the less noise is required to produce alternations. Within the switching domain different combinations of *T*_*a*_ and *σ*_*a*_ yield different distributions and means of percept durations. Our model reproduces the experimental data when *T*_*a*_, *σ*_*a*_ are taken from a restricted parameter region. With *T*_*a*_ constant across conditions we captured the observed trend of mean durations although some values were off. With fine-tuning of *T*_*a*_ across conditions (but *σ*_*a*_ constant) we match the observed duration distribution shapes and means. This approach is analogous to obtaining the proper balance between noise and adaptation necessary for alternations in other models for bistable perception [22, 33]. Noteworthy, our model shows switching behavior when tuned in other parameter regimes, including with *T*_*a*_>1. However, in such a drift-dominated regime although noise is not needed for alternations, we found that matching the statistical features and behavioral trends required a substantially higher (unacceptable) noise level (not shown here).

### Comparison with other models

Barniv and Nelken (2015) and Cao et al (2016) also modeled auditory bistable perception as evidence accumulation based. The former’s model used Bayesian assignments of *B*-tones to either the same class as *A*-tones (integration) or to a different class (segregation). Its noise-free version shows periodic alternations, as does our system for *T*_*a*_>1, but the dynamics do not reset. Instead, our accumulators undergo discontinuous resetting after each switch. Most importantly, in contrast to [5], the parameters in our model are interpretable, functionally if not physiologically. Cao et al. formulated a stochastic accumulator that reproduced (like ours) several scaling properties of bistable behavior but without a description of switching and of, possibly asymmetric, alternations. Our work differs from both models by incorporating directly A1-spiking data as input. In this sense it is more akin to [6], a literal competition model. Notably, our approach predicts that neuronal computation for percept representation and evidence accumulation takes place beyond A1; its input (activity from A1 devoid of switch-dynamics) implicitly includes inhibition, adaptation and noise that occur within A1 and preceding stages.

Dynamic competition models commonly include two or more units representing response patterns associated with different percepts, and share mechanistic features of mutual inhibition, adaptation, and noise [6, 14]. In our two-process model only one percept is currently dominant thereby realizing mutual exclusivity. However, inhibition is not explicitly invoked; rather, our model performs as if a firm choice is made at the switch time, further accumulation of evidence in favor of the fresh percept is prevented; the in-favor accumulator is reset and targeted to a low value, *T*_*f*_, despite continued incoming evidence from SL.

In oscillator-based alternations, switching may be determined by strongly rising adaptation in the dominant unit, leading to “release” from inhibition, or by stronger recovery from adaptation in the suppressed unit, leading to “escape” from inhibition [34–37]. Our model has no explicit adaptation variable as negative feedback. However, the accumulation of evidence against the current percept may be viewed as recovery of salience of the non-dominant percept. The rise and eventual take-over of dominance is therefore analogous to the escape from suppression in competition models.

In such models if adaptation is weak, changes in dominance may be represented by noise-driven switches between stable states in attractor state dynamics [22, 23]. These insights motivated our choice of an evidence-against accumulator that saturates just-subthreshold. Dominance durations are longer with reduced noise, and no switches occur in the noise-free idealization. Further, gamma-like duration distributions are more robustly obtainable with this mechanism: rise to saturation and wait for switch-favoring fluctuations to induce a switch. Satisfactory results are also obtainable with *T*_*a*_>1, but if *T*_*a*_ exceeds threshold by too much, acceptable duration statistics seemed to require strong noise, and accumulator time courses were noise-dominated.

Bistable perception for ambiguous visual displays was modeled by [38] as a continuous time accumulation of binary (bistable) units becoming active with state-dependent transition rates between the active and inactive states. Our modeling shares a key feature: saturation to a level that strongly affects the percept durations; saturation near/below threshold underlies escape-like dynamics with gamma-like duration distributions. Distinguishing from [38], our model is event-based (discrete-time) with stimuli-induced positive increments and additive zero-mean noise that allow positive/negative increments, not a Markov model. It is applied directly to the neural data and includes saturation with noise-driven attractor dynamics as in competition models.

### Limitations, extensions, predictions

Reports on triplet-streaming are conflicted about correlations between successive *I, S* durations, showing either statistical independence [2] or small positive correlations [5]. In our model both accumulators are reset after a switch approximately to target *T*_*f*_ so correlation between successive percepts is weak. However, we could likely match the reported correlations [5] by changing the resetting to generate continuous dynamics of accumulators.

Alternations between percepts are generated by the evidence that accumulates against the current percept. Its dynamics depends primarily on the distance to target *T*_*a*_ and on input *p* from the sampler layer, with *T*_*a*_ assumed relatively constant across conditions. Alternatively, one might choose *T*_*a*_ as a *DF* -dependent parameter and keep *p* unchanged. Such an approach suggests an interpretation of the target, with nearness to threshold, reflecting a combination of condition-dependent input and inhibition, and possibly excitation (in-line with a population separation hypothesis [7]). Then *p*, as constant and independent of *DF*, can be viewed as rate of recovery from adaptation. However, to establish a derivable connection between *T*_*a*_ and the experimental condition and A1 spiking activity presents challenges.

In our model the number of A1 neurons that are sampled by each SL-unit is much lower than the number of recorded units used in the signal detection approach in [1] (*N*_*in*_=5 vs 91 cortical neurons). We found that the granularity of sampling A1 by a unit in the sampler layer is important in order to preserve sufficient variability in the averaged spike count over trials and thereby obtain graded BUFs across different DF conditions. Perhaps the constraint on *N*_*in*_ derives from our assumption of statistically independent A1 neurons. As shown by [27], trial-to-trial variability in spike counts for *N*_*in*_ small, if spikes are statistically independent, is equivalent to the variability over a much higher number of A1 neurons if correlations exist within the pool. We did not have access to the original spike times from [1] to verify this hypothesis; we only extracted mean spike counts from the published data. However, this observation is supported by a subsequent study by Micheyl et al [12] and could be explored in future simulations; when spike counts from a subset of 30 cortical neurons (or even just one neuron) out of 91 were analyzed with the signal-detection model, the resulting neuronal-based BUFs were less widely spread across conditions, matching the graded behavioral BUFs from a different subject pool (see [12, Fig.5], compare with Figs 3 and 8 for our model).

Our model could be extended to mimic the transient behavior of buildup by relaxing the constraints on initial conditions and treating the baseline as *DF* -dependent. Two hypotheses may be tested: that integration emerges with first percept probability as in the behavioral data and that early adaptation of A1-responses accounts for longer first, than subsequent, *I*-durations [39].

With minimal modifications to our model we could test for behavior at other *DF* values or for dependence on presentation rate. Assuming lower target-against levels *T*_*a*_ for faster presentations, we predict at constant *DF* similar mean *I*-durations but longer *S*-durations, and higher probability of segregation [6]. With increased presentation rate mean spike counts for *B*-tones will decrease [8] and lead to lower vote counts *p*_*I*_ and lower effective accumulation rate, *T*_*a*_*p*_*I*_. Although *p*_*S*_ (=1-*p*_*I*_) would increase, the increase would be compensated by the decreased *T*_*a*_ leaving *T*_*a*_*p*_*S*_ relatively unchanged. To conclude, we propose an evidence accumulation model for auditory bistable perception with neurally-plausible mechanisms that accounts for statistics of behavioral data. In principle, it could be extended to study dynamics induced by transient perturbations (deviants/distractors; [39]) or associated with multiple percepts [14]; implementations of such generalizations remain as open topics for future research.

## Methods

### Experimental design and statistical analyses

#### Participants

Fifteen human subjects with normal hearing (8 female and 7 male; ages 18-45 yrs.; median 22 yrs.) were included in the behavioral study. They listened to sequences of repeating *ABA*_ triplets and were instructed to continuously report their ongoing percept by selectively pressing one of two different buttons on a keypad. Subjects began reporting their percept typically 2 s after stimulus onset as integration (*I*; a single, coherent stream, the galloping pattern *ABA*_*ABA*_) or segregation (*S*; two simultaneous distinct streams *A*_*A*_*A*_*A*_ and _*B*___*B*_); Fig 1 A-B.

#### Stimuli

Stimuli were 30-s long sequences of triplets *ABA*_ that consisted of alternating high (*A*) and low (*B*) pure tones gated with 10 ms raised cosine ramps and followed by a 125 ms silent pause “_”; Fig 1A. In total, triplets were 500-ms in duration and were repeated 60 times per trial. Tones were separated in frequency by *DF* semitones chosen from three conditions (*DF* = 3, 5, 7) with each condition being presented five times per experimental block. To prevent habituation to a certain frequency, for each *DF* the tones were generated by roving through variants of frequencies taken 0, ±1 or ±2 semitones apart from their geometric mean (middle pair in the list below); see also [4]. Frequencies (*f*_*A*_, *f*_*B*_), in Hz, were chosen as: (494, 415), (523, 440), (554, 466), (587, 494) or (622, 523) Hz at *DF* =3; (523, 392), (554, 415), (587, 440), (622, 466), (659, 494) at *DF* =5; and (554, 370), (587, 392), (622, 415), (659, 440), (698, 466) at *DF* =7. Stimuli were digitally generated in Matlab at 48 kHz sampling rate and were delivered through earphones in a soundproof isolated room. Subjects had the sound volume adjusted to their comfortable hearing level.

#### Experimental protocol

Each subject performed 9 experimental blocks. Each *DF* condition was randomly presented 5 times per experimental block, using different combinations of frequencies for tones *A* and *B*, without repetition (see *Stimuli*). A Latin square design was used to determine the order of presentation of each condition in each block. This resulted in group data with 675 30-s trials for each of three *DF* values. The frequency separation values (*DF* = 3, 5, 7) were chosen to fall within the range of ambiguity of the van Noorden diagram in which listeners can perceive both integration and segregation [10, 40]. All subjects underwent a training session in which they were given verbal explanations and auditory illustrations of the two possible percepts, and they practiced distinguishing between them. Then, during the recording session, listeners were instructed to press and hold one key on a keypad when they perceived stimuli as *I*, and to release it while pressing another key when they perceived stimuli as *S*, and so on. They were encouraged to respond as soon as they heard the change in percept. The key-response data were converted to binary vectors with value 0 assigned to *I* (and to the latency period defined as the time before identification of either percept) and 1 to *S*, for further analysis. Experiments were performed in a dedicated soundproof booth in the Human Brain Research Laboratory, Neurosurgery Department at The University of Iowa. Written informed consent was obtained from all subjects. Research protocols were approved by the University of Iowa Institutional Review Board.

#### Build-up functions and the latency period

The time course of *S* percept after stimulus onset was computed from key-pressed data, 0 for *I* and 1 for *S*. Those were sampled at 1 ms to create vectors of binary values corresponding to appropriate percept type at a particular time instance. At each *DF* condition, binary vectors were averaged across 675 trials to obtain the build-up function of *S*; bootstrapping was also used to compute the 95% confidence interval (CI) around the mean (Fig 1 B-C). The time course of *I* was computed with the same procedure but over key-pressed data labeled as 1 for *I* and 0 for *S*. At a particular time *t*, the proportion of trials classified as *I, S* or neither (during the latency period/the first few seconds after stimulus onset) were *p*_*int*_(*t*), *p*_*seg*_(*t*) and *p*_*Lat*_(*t*), and summed up to *p*_*int*_(*t*) + *p*_*seg*_(*t*) + *p*_*Lat*_(*t*) = 1. The first percepts were typically *I*. However, for larger values of tone frequency separation, subjects tended to report *S* as first percept more often which led to an increase in *p*_*seg*_(*t*).

#### First durations and subsequent durations

All complete percept durations across trials and conditions were included in the behavioral analysis. Unfinished percepts and button presses recorded after the end of stimulus presentation were discarded. For each *DF* = 3, 5, 7, the statistics was evaluated over four subsets of data, separately: first *I*, first *S*, subsequent *I* and subsequent *S*. For each *DF* and each of these four percept types, the mean dominance duration was computed in two steps: first, it was computed per subject, say *µ*_*i*_ for subject *i* = 1, *…*, 15; then the mean duration *µ* of the group data was defined (and reported) as the unweighted average across all subjects, *µ* = (*µ*_1_ + *µ*_2_ + *· · ·* + *µ*_15_)*/*15 (e.g. Fig 1C; mean *µ* is shown for first duration distributions at *DF* =5). By this approach, any potential bias of the calculation towards fast switchers who might contribute more durations to the pool and concurrently spend less time in a particular percept, was mitigated. Error bars at 95% CI of the mean were also determined; error bars corresponded to 1.96 SE; standard error 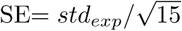 was computed from the standard deviation *std*_*exp*_ over the group of means *µ*_*i*_. For analysis of grouped data we used subject-specific normalized (by individual subject mean) percept durations as follows: at each *DF* condition and each percept type (first/subsequent, *I*/*S*), any raw percept duration *D* of subject *i* was normalized by the corresponding mean *µ*_*i*_ to 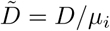. Histograms of normalized phase durations 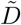 for each *DF* condition and percept type were computed and fit by gamma distributions with density functions

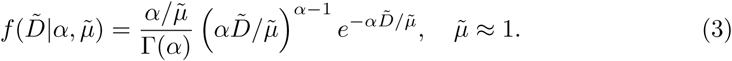

Mean 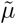 was well-fit to 1 due to normalization. Then the coefficient of variation 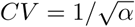 and the skewness 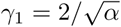 depended exclusively on the shape parameter *α*. If *α* was large then (3) was equivalent to a normal distribution. If *α* ≈ 1 then (3) was equivalent to an exponential distribution. On the other hand, for *α* ≈ 2 (as observed for first durations in behavioral data) and *α* ≈ 2.6 (as observed for subsequent durations), the distributions satisfied the scaling property *γ*_1_ = 2*CV* with *CV* ≈ 0.7 and *CV* ≈ 0.6 respectively. The latter case was similar to the findings of [24] that described the statistics of percept durations for other examples of perceptual bistability.

#### Distribution testing of phase durations

The fitting of the experimental (and numerical) data was obtained by calculating the values of *α* and 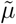 with the Maximum Likelihood Estimation (MLE) algorithm. The goal was to determine *α* and 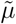 that yielded the maximum product п _*k*_ *y*_*k*_ of all *y*_*k*_ gamma-likelihood values of the normalized percepts 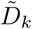 counted by index *k* for each run of the experiment. This was equivalent to maximizing the log-likelihood 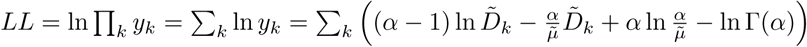 based on formula (3). The optimization of *LL* was implemented numerically with MATLAB function fminsearch. Distribution testing on normalized durations was done by statistical bootstrapping. We generated 10000 bootstrapping sets of gamma distributions with fitted parameters *α* and 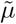 and constructed the distribution of maximum log likelihood values for those sets. The test statistics *LL* was compared to this distribution to obtain the probability of log likelihood to be less than *LL* (the *p*-value). The normalized durations were well fit by a gamma distribution with the optimal values *α* and 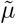 (as null hypothesis) at significance level 0.05 if *p*-value ≥ 0.05.

#### Statistical analysis of model-generated data

The histograms of first and subsequent durations *I* and *S* in trials generated by the model (see below), and their fitting by gamma distributions, were computed in a similar manner as for the experimental data. Likewise, build-up functions for the model were constructed as those for the behavioral data.

### The evidence accumulation model

Our proposed EVA model is a feedforward network of 3 layers: the input layer of spiking units, the sampler layer of binary response units, and the accumulation layer of two accumulators. The time-unit of the model is discrete and defined as the position of the triplet in the auditory sequence. Every *DF* -dependent numerical simulation of the EVA model consisted of *N*_*tr*_=675 repetitions (trials) to mimic the setup from the behavioral experiment. The trials were then used to generate the statistics of the percept durations in terms of mean values, shape of distributions, and buildup functions. Finally, in order to test for the model’s robustness in the presence of noise (not for sensitivity to model parameter values), this numerical procedure was run 100 times for each *DF* = 3, 5, 7 condition separately, and the results were characterized by averaged values and their 95% CI.

#### Input layer (IL)

The IL-units were assumed to be tone-*A* selective neurons from primary auditory cortex (A1) as described in [1]. The averaged (over trials) spike counts *m*_*t,DF*_ of the IL-units were derived from data (see section *Data-driven parameters for EVA model*) and depended on the position *t* of the triplet in the *ABA*_ sequence (*t*=1, *…*, 60 for a 60-triplet long stimulus; 30 s in duration) and on the semitone difference *DF*. The model was simplified by focusing only on the spike counts during the *B*-tone presentation at *A*-tone selective neurons in A1. As reported by multi-unit recordings in monkeys, such A1-neurons adapted strongly and rapidly during presentation of triplet-repeating auditory sequences [1, total of 91 neurons]. Temporal correlations between the means of an A1-neuron from triplet to triplet were captured in the model by the trend of *m*_*t,DF*_ that decreased exponentially with *t* (Fig 4B). Trial-to-trial variability of the dynamics of IL units as well as unit variability in IL during a single trial were implemented using Poisson point processes. (We used this approach because we could extract mean spike counts from published data of [1] but did not have access to the original spike times.) For an A1-neuron with mean spike count *λ*= *m*_*t,DF*_ we supposed that its instantaneous spike count *k* (*k* = 0, 1, 2, *…*) at triplet *t* and condition *DF*, was randomly generated from a Poisson distribution with probability *P* (*X* = *k*) = *λ*^*k*^ *e*^−*λ*^*/k*!. The Poisson spike counts are generated independently for each neuronal unit in IL, each triplet and each semitone difference condition. Note that the model could be generalized by assuming neuronal heterogeneity, with averaged spike counts 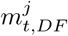 at neuron *j* chosen from a normal distribution *𝒩* (*m*_*t,DF*_, *s*_*t,DF*_) of mean *m*_*t,DF*_ and standard deviation *s*_*t,DF*_ derived from [1]. However, the impact of heterogeneity on the model’s outcome would be negligible given that EVA model was formulated to use mean spike counts over IL neuronal pools as input rather than mean spike counts of individual neurons (see below).

#### Sampler layer (SL)

The SL-units were tasked with summating and classifying spike counts from subsets of *N*_*in*_ IL units (A1-neurons). Consider that a trial of length *N*_*t*_ (*N*_*t*_=60 triplets) during a certain *DF* condition was simulated by EVA model: each sampler summed the input of a pool of *N*_*in*_ neuronal units from IL; weighted by *N*_*in*_, this gave the mean spike count for *B*-tones of the corresponding pool of *A*-tone selective A1-neurons, 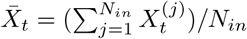 for triplet *t*; then 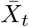 was compared to a *DF* -independent, fixed, neuronal threshold *C*_*th*_. If the averaged spike count was large 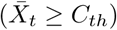 then the subset activity was high and the pool was assumed to support, for this triplet, the integration percept *I*. The sampler’s response at triplet *t* was classified as “1”. If the averaged spike count was small 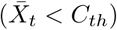, the subset activity was low and the IL-pool was said to support the segregation percept *S*. The sampler’s response was classified as “0” (Fig 4A). Therefore, at each triplet *t*, each sampler behaved like a biased coin being flipped with probabilities *p*_0;*t,DF*_ and *p*_1;*t,DF*_ over the binary probability space of 0 and 1. Since neuronal units in each IL-pool followed independent Poisson distributions of parameters *m*_*t,DF*_, the pool itself was also a Poisson process defined by the product *N*_*in*_ *m*_*t,DF*_. Then, the samplers were binary signal detectors with probabilities

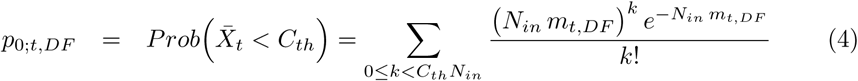

and *p*_1;*t,DF*_ = 1 − *p*_0;*t,DF*_ calculated over 675 repetitions of the model in order to maintain similarities to the behavioral experiment, and with expected value and variance 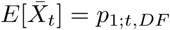and 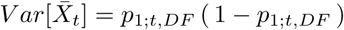.

Three *DF* -independent parameters were associated with SL: *N*_*in*_, the number of A1 inputs to a sampler unit; *C*_*th*_, the neuronal counting threshold that categorizes ensemble activity in A1 as high (class 1) or low (class 0); and *N*_*sl*_, the number of neuronal units in SL. The values of *N*_*in*_ and *C*_*th*_ were obtained by least-squares fit between probabilities *p*_0;*t,DF*_ and the “asymptotic” levels 0.45, 0.6, 0.65 of the psychometric buildup functions (last 15 seconds of trial duration) for all *DF* = 3, 5, 7 (Fig 8B; also section *Data-driven parameters for EVA model*). The psychometric buildup represented the fraction (over the trials) of the segregation percept *S* reported by all subjects at each time point and *DF* during the 30-s long trial. Through this optimization procedure (Fig 8B; optimal values obtained for *N*_*in*_ = 5, *C*_*th*_ = 4.21) probabilities *p*_0;*t,DF*_ of a sampler to be in a state that supported percept *S* were estimated – at least for triplets several seconds from the stimulus onset – as

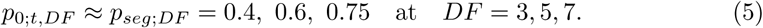

The inclusion of the sampler layer in the model (*N*_*sl*_ > 1) ensured neuronal granularity that was found by other studies to be a suitable substrate for perceptual representations [24]. In particular, at each triplet *t*, some of the *N*_*sl*_ samplers were in class 0 showing low A1 spiking and thereby associated with segregation [1, 7] while others were in class 1 supporting integration. The percentages *p*_*S*_ (*t*) and *p*_*I*_ (*t*) = 1 − *p*_*S*_ (*t*) of such samplers were taken herein as stochastic (over trials) output of SL (Fig 4A).

#### Accumulation layer (ACC)

The accumulation layer consisted of two units whose dynamic states *x*_*I*_ and *x*_*S*_ changed from triplet to triplet according to Eqs

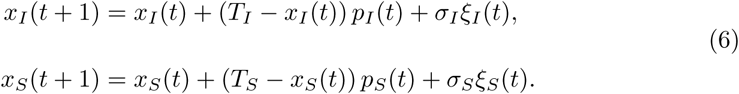

The accumulator for *x*_*I*_ gathered evidence that favored integration (through input *p*_*I*_ (*t*) from SL) while the accumulator for *x*_*S*_ gathered evidence that favored segregation (through input *p*_*S*_ (*t*) from SL). Importantly, their states were influenced by the perceptual context as well (Fig 4C, solid lines in blue and red illustrated traces for *x*_*I*_ and *x*_*S*_; the background color showed the percept’s type, blue for *I* and red for *S*). In particular, if *segregation* was the current dominant percept then *x*_*I*_ accumulated evidence *against* segregation and aimed to reach target *T*_*I*_ =*T*_*a*_; simultaneously, *x*_*S*_ accumulated evidence *for* the current percept and it drifted instead towards target *T*_*S*_ =*T*_*f*_. Discrete-time Gaussian white noise processes *σ*_*I*_ *ξ*_*I*_ (*t*), *σ*_*S*_*ξ*_*S*_ (*t*) with zero mean and standard deviations *σ*_*I*_ =*σ*_*a*_ and *σ*_*S*_ =*σ*_*f*_, interacted with the stochastic inputs from SL to produce certain levels of fluctuations. The additive stochastic terms in ACC were target-dependent with *σ*_*a*_ > *σ*_*f*_ for *T*_*a*_ > *T*_*f*_ (Fig 4C). On the contrary, if the current percept was integration then *x*_*I*_ accumulated evidence for *I* and approached *T*_*I*_ =*T*_*f*_ while *x*_*S*_ accumulated evidence against *I* and approached *T*_*S*_ =*T*_*a*_. The level of local noise was adjusted accordingly to values *σ*_*I*_ =*σ*_*f*_ and *σ*_*S*_ =*σ*_*a*_.

#### Switches and resetting conditions

In the experiment, subjects identified the dominant percept by pressing a certain button on the keypad. Equivalently, in EVA model, the switch from one dominant percept to the next occurred when either *x*_*I*_ (*t*) or *x*_*S*_ (*t*) crossed the ACC threshold set to 1. Herein, the alternations were initiated by the accumulator that observed how many samplers in SL opposed the current percept at each triplet *t*. Its trace was attracted to target *T*_*a*_ that lay near the threshold, then was pushed across the threshold by the noise of strength *σ*_*a*_. In the meantime, the accumulator in favor of the current percept hovered around *T*_*f*_ with fluctuations set by *σ*_*f*_. For example during percept *I*, accumulator *x*_*S*_ was the first to reach threshold 1 producing a switch to percept *S*; then, during *S, x*_*I*_ reached threshold 1 leading to another switch to subsequent percept *I*, and so on (Fig 4C). At every change in percept, the accumulators were reset to new levels 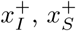 These were defined as 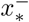 where 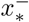 was the value that the evidence-for accumulator *x*_*_ (* = *I, S* during current percept *I, S* respectively) reached just before the switch.

The simulations took into account the proportion of first percepts reported as segregation at each *DF* during the behavioral experiment as well as the latency period during each trial. The initial conditions of the accumulators were set to a *DF* -independent baseline value *b* and kept constant during the entire latency period (calculated in length of *T*_*Lat*_ triplets) of any given trial. We defined *x*_*I*_ (*t*) = *x*_*S*_ (*t*) = *b* for all triplets *t* between 1 and *T*_*Lat*_; then at *t* = *T*_*Lat*_ the type of the current first percept, *I* or *S*, was imported from the behavioral data; the dynamics of the accumulators for*t* ≥ *T*_*Lat*_ were then determined according to Eqs (6) and the associated reset conditions.

The choice of parameters (*σ*_*f*_ small) and of reset conditions (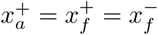 where *x*_*a*_, *x*_*f*_ are accumulators “against” and “for” the dominant percept) ensured that the switch was triggered by the dynamics of *x*_*a*_. Rare events when *x*_*f*_ might have crossed the threshold ahead of *x*_*a*_ were disregarded. Another possible implementation of resetting would allow for correlations between consecutive percepts; it could depend on each accumulator state just before a switch, a simple interchange of roles such that ACC variables remained continuous, 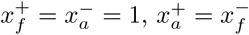 (not shown in this paper).

### Model analysis

#### Parameter values used in model simulations

All figures for the full EVA model (Figs 1, 4 – 8, and S1 Fig – S3 Fig, S5 Fig), were generated with the following parameter values:

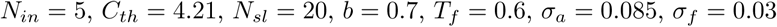

and decision threshold *θ* = 1, unless otherwise stated in their caption. (For parameter values *m*_*t,DF*_ associated with IL, see *Data-driven parameters for EVA model*.) Target *T*_*a*_ was initially chosen equal to 0.9 and then was fine-tuned to best fit the mean dominance durations of the first and subsequent integration and segregation percepts from the behavioral data – within ±10% standard error (SE) of the experimental mean values (Figs 5 and 6; also S5 Fig). Its values changed with *DF*, and with classification (first or subsequent) and type (*I* or *S*) of the percept. We used the notation *T*_*aI*1_ for “target against integration, first percept”, *T*_*aS*1_ for “target against segregation, first percept”, *T*_*aI*2_ for “target against integration, subsequent percepts”, and *T*_*aS*2_ for “target against segregation, subsequent percepts”, respectively. Therefore, in the model, whenever acting as target-against, *T*_*S*_ = *T*_*aI*1_ or *T*_*aI*2_ while *T*_*I*_ = *T*_*aS*1_ or *T*_*aS*2_. In simulations we used the following values (see red diamonds in Fig 7):

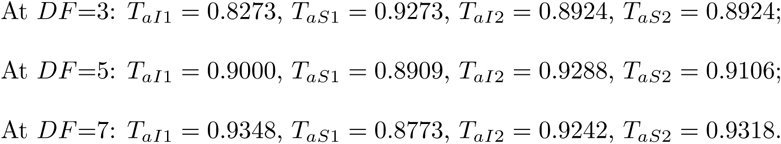

#### Selection of target-against *T*_*a*_ values

The mean duration *µ* obtained by numerical simulations of EVA model was compared to its behavioral counterpart *µ*_*exp*_ for first and subsequent *I, S* percepts, and each *DF* = 3, 5, 7. Behavioral results from 15 subjects were characterized by group mean data *µ*_*exp*_ and 95% CI with CI corresponding to 1.96 SE (Fig 5A and Fig 6A, lower panel). Then the model was considered to provide a good approximation of the experimental data if *µ* belonged to a narrow band within 1 SE from *µ*_*exp*_ (Fig 7, green region and black dots). This was equivalent to the relative error 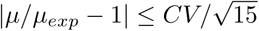where *CV* = *std*_*exp*_*/µ*_*exp*_ was the coefficient of variation computed over the group of subjects. Parameters *T*_*a*_, *σ*_*a*_ used for the numerical simulations of EVA model (Fig 7, red diamond) were chosen as follows: *σ*_*a*_ = 0.085 was kept fixed while values of *T*_*a*_ were determined by restricting the error magnitude to only 10% SE (i.e. *|µ* − *µ*_*exp*_*| ≤* 0.1 SE); then, among the latter set we selected the value *T*_*a*_ that yielded the least error in shape of the gamma-fit distributions (see Eq (3)).

#### Statistical properties of SL-activation

At any fixed triplet position *t* in the *ABA*_ sequence presentation, each of the *N*_*sl*_ samplers was equivalent to an independent Bernoulli process (during repeated trials) with probability of success *p*_1;*t,DF*_ and probability of failure *p*_0;*t,DF*_ = 1 − *p*_1;*t,DF*_. Likewise, the state of SL described a binomial process equivalent to the random variable, over trials, *N*_*sl*_ *p*_*I*_ (*t*) where *p*_*I*_ (*t*) represented the percentage of samplers in class 1 that favored integration at triplet *t*. The stochastic process had mean *N*_*sl*_ *p*_1;*t,DF*_ and variance *N*_*sl*_ (1 − *p*_1;*t,DF*_) *p*_1;*t,DF*_. As a result, the first two moments of the output *p*_*S*_ (*t*) and *p*_*I*_ (*t*) of SL were well-approximated, for sufficiently large triplet-indexes(t ≥ 30; see Eq (5) and Fig 8B, 2nd panel), by

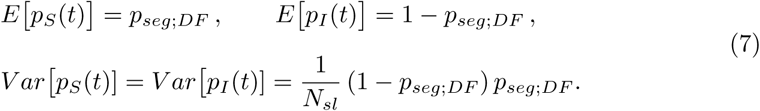

In particular, for large *N*_*sl*_ the variance of *p*_*S*_ (*t*) and *p*_*I*_ (*t*) became negligible while their means remained unchanged.

#### Selection of noise level *σ*_*a*_

Our EVA model features accumulation that could saturate, given that target-against *T*_*a*_ was assumed to be subthreshold (*T*_*a*_ < 1). Hence the noise level *σ*_*a*_ has to be sufficiently large in order for the trajectory of the accumulator drifting towards *T*_*a*_ to reach the ACC-threshold. A theoretical lower-bound estimate for *σ*_*a*_ was obtained by assuming *N*_*sl*_ large and focusing only on the properties of the subsequent percept durations. Under such assumptions, *p*_*S*_ (*t*) and *p*_*I*_ (*t*) were approximately constant as demonstrated by Eqs (5) and (7). Then, after each switch, both accumulators satisfied an equation of the form *X*_*n*+1_ = *X*_*n*_ + (*T* − *X*_*n*_)*p* + *ε*_*n*+1_ with *X*_0_ ≈ *T*_*f*_; *T, σ* taken as either *T*_*a*_, *σ*_*a*_ or *T*_*f*_, *σ*_*f*_; and independent random variables *ε*_*n*+1_ ∼ *N* (0, *σ*^2^); see also Eq 1 in Section B.1. Such an equation describes a stationary first order autoregressive model with parameter *λ*=1-*p* [41]. *Therefore, at the n*th triplet during a percept immediately following a switch, the states of the accumulators followed a normal distribution with mean *E*[*X*_*n*_]=*T* -(*T* -*T*_*f*_)*λ*^*n*^ and variance *V ar*[*X*_*n*_]=*σ*^2^(1 − *λ*^2*n*^)*/*(1 − *λ*^2^). In particular, at the *n*th triplet during integration, the mean and variance of *x*_*S*_ (*x*_*I*_) that accumulated evidence against (for) the percept were *E*[*x*_*S*_] = *E*_*a*_, *V ar*[*x*_*S*_] = *V*_*a*_, *E*[*x*_*I*_] = *E*_*f*_, *V ar*[*x*_*I*_] = *V*_*f*_ where

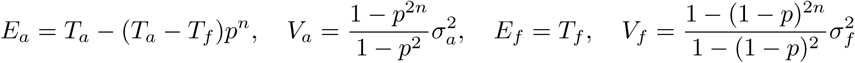

with *p* = 1 − *p*_*seg;DF*_. Then at the nth triplet during segregation they were *E*[*x*_*I*_] = *E*_*a*_, *Var*[*x*_*I*_] = *V*_*a*_, *E*[*x*_*S*_] = *E*_*f*_, *Var*[*x*_*S*_] = *V*_*f*_ with *E*_*a*_, *V*_*a*_, *E*_*f*_, *V*_*f*_ defined as above but computed with *p* = *p*_*seg*;*DF*_. Given that 3 times the standard deviation from the mean accounts for 99.73% of values in a normal distribution, if *σ*_*a*_ was too small the accumulators could cross the threshold 1 only with very small probabillity. From the calculation above, a lower bound for *σ*_*a*_ in the model was estimated at *σ*_*a*_ > *σ*_*a,min*_ with 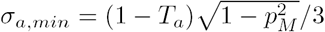, where *p*_*M*_ was the maximum of *p*_*seg*;*DF*_ and 1 − *p*_*seg*;*DF*_ for all *DF* = 3, 5, 7. For example, if *T*_*a*_ = 0.9 then a necessary condition for switching was *σ*_*a*_ > 0.022.

#### Statistical properties of ACC-activation

As explained in the previous section, for small *σ*_*a*_ the accumulators in the EVA model could not cross threshold 1 (Fig 7; na, gray region). Numerical simulations showed that when *σ*_*a*_ increased to the right of curve *σ*_*a*_ = *σ*_*a,min*_ in the (*σ*_*a*_, *T*_*a*_)-plane, alternations between percepts occurred and the dominant durations were distributed according to: normal distributions at *σ*_*a*_ small (the parameter *α* in Eq (3) was very large) or to exponential distributions at *σ*_*a*_ large (*α* in (3) was near 1); see Fig 7, regions labeled “N” and “E”, respectively. At intermediate values *σ*_*a*_, the distributions were gamma-like with shape close to that found experimentally (Fig 7, region labeled “G” between the two white curves). In the latter case, parameter *α* in (3) was either near 2 (for first percept durations) or near 2.6 (for subsequent durations), and it differed from *α*_*exp*_ by relative error up to 20%, *|α/α*_*exp*_ − 1*| ≤* 0.2 (except for first and subsequent *I* at *DF* =7 at which the range for *α* was extremely narrow and we allowed for a larger error, up to 30% instead). The range for *σ*_*a*_ that led to gamma-like distributions varied slightly with *N*_*sl*_ with the biggest difference being identified at *N*_*sl*_ = 1; see S3 Fig.

#### Data-driven parameters for EVA model

Parameter mean values *m*_*t,DF*_ were used to generate Poisson spike counts of IL-units at each triplet *t* (*t* = 1, 2, *…*, 60) in the *ABA*_ sequence and for each semitone difference *DF* =3, 5, 7; see section *Input Layer* and Eq (4). They were derived from multi-unit spiking neural data recorded from macaque monkey primary auditory cortex A1 by [1], using a procedure that combined exponential fitting with numerical interpolation. First, mean spike counts *m*_*t,DF*_ at A-tone selective neurons in A1 during presentation of tones *A, B* and *A* in *ABA*_ were extracted from [1] for each triplet *t ≤* 20 in the sequence and for each *DF* =1, 3, 6, 9; See scatter points in [1, Fig.3]; also S4 Fig. The mean spike counts at each tone decreased from value *m*_1,*DF*_ measured at the first triplet to some level 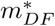 at which they stabilized after a few seconds since stimulus onset. They were fitted by functions

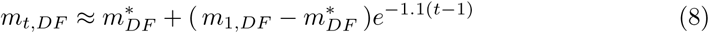

with parameters 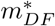 chosen to minimize the least-squares error between the extracted mean spike counts *m*_*t,DF*_ and the corresponding exponential curve (S4 Fig, solid curves). In particular, significant differences in mean spike counts at different *DF* values were observed only during tone-*B* presentation with fitting (8) achieved for parameter values *m*_1,1_ = 7.25, *m*_1,3_ = 6.25, *m*_1,6_ = 6, *m*_1,9_ = 5.25 (according to data from [1]) and 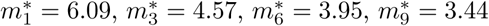 respectively. Secondly, simulations of EVA model were performed for *DF* = 3, 5, 7 instead of 1, 3, 6, 9, and for a total of 60 rather than 20 triplets. We implemented these constraints in two steps: for *DF* =3 we chose the mean spike counts *m*_*t,DF*_ as in [1] for *t ≤* 20 and as 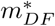 for *t* > 20. Then for *DF* =5 and *DF* =7 and each triplet *t* we defined the mean spike counts by interpolation using the power function 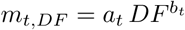 whose coefficients *a*_*t*_, *b*_*t*_ satisfied the least-squares fit between this curve and the points (*DF, m*_*t,DF*_) defined by for all *DF* =1, 3, 6, 9 at any fixed *t*.

The mean spike counts for *B*-tones of any pool of *A*-tone selective IL neuronal units, as well as the standard error to the mean, were computed from simulation of Poisson processes with parameter *m*_*t,DF*_ while varying *N*_*in*_ (Fig 8A). Then a threshold value *C*_*th*_ was chosen to minimize the squared differences error between the model-based probabilities (4) of a sampler to support the segregation percept for all *DF* =3, 5, 7 and the behavioral buildup functions, applied to the last 30 triplets (15 seconds) of the stimulus (Fig 8B). The pair of parameter values *N*_*in*_ = 5, *C*_*th*_ = 4.21 that generated the minimum error was then used in numerical simulations of the EVA model.

### EVA model versus classical drift-diffusion models

To gain some intuition about the accumulation process in our EVA model and about the timing of switch events, we considered an approximation of the stochastic Eqs (6) in continuous time. For that, we assumed the drift in (6) to be constant and neglected its dependence on activity *x*. Then percept durations corresponded to the first-passage time of the ACC-unit that accumulated evidence against the current percept. Its equation could be interpreted as the constant-drift continuous-time diffusion model (DDM) *dx* = *γ*_*a*_*dt* + *σ*_*a*_ *dW*_*t*_ with positive drift rate *γ*_*a*_, noise amplitude *σ*_*a*_, Gaussian white noise *dW*_*t*_, and decision threshold *θ* = 1. In this DDM, the likelihood of first-passage at time *t* follows an inverse Gaussian distribution [28] with density function 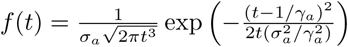 and mean 1*/γ*, variance 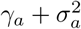, and coefficient of variation 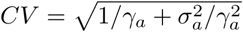. Moreover, the inverse Gaussian resembles a gamma distribution for large *CV* but converges to a normal distribution as *σ*_*a*_ decreased in relation to drift rate *γ*_*a*_ [28]. To some extent, the dynamics of the discrete-time ACC (6) share similarities with DDM above. Numerical simulations of our EVA model showed that gamma-like distributions of percept durations were possible only for *σ*_*a*_ chosen in a restricted parameter range, given fixed targets *T*_*a*_ and *T*_*f*_ (see section *Statistical properties of ACC-activation*). Outside this range, percept durations followed either normal distributions (for lower values of *σ*_*a*_) or exponential distributions (for larger values of *σ*_*a*_). However, the accumulation process in the EVA model is different than in the DDM for several reasons: Eqs (6) are discrete-time drift diffusion models; they include leakage; their deterministic version admits bistable non-oscillatory solutions (no threshold crossing); and the input drive from SL is itself stochastic with fluctuations described by (7).

## Acknowledgments

The authors thank Xiayi Wang, Haiming Chen and Kirill V. Nourski for help with data acquisition, and Dan Tranchina for suggestions on data analysis.

## Supporting information

**S1 Fig.**
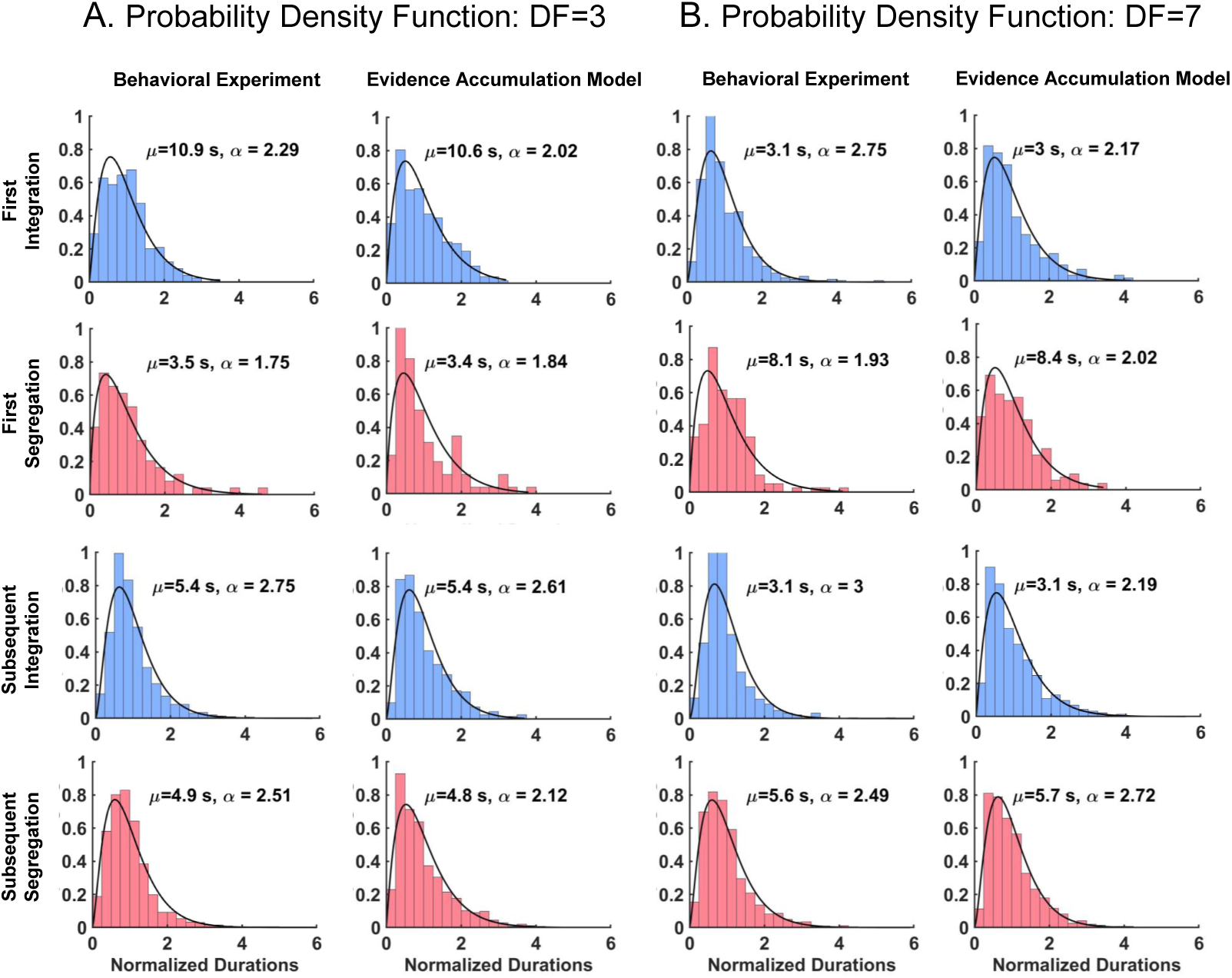
The evidence accumulation (EVA) model captures experimental mean duration (*µ*) and shape of gamma-like distributions (*α*) for first and subsequent percept durations at other *DF* values. Distributions of normalized phase durations are shown for A: *DF* =3 and B: *DF* =7. They are obtained from numerical simulations of the EVA model (columns 2,4) and compared to those derived from behavioral data (columns 1,3).

**S2 Fig.**
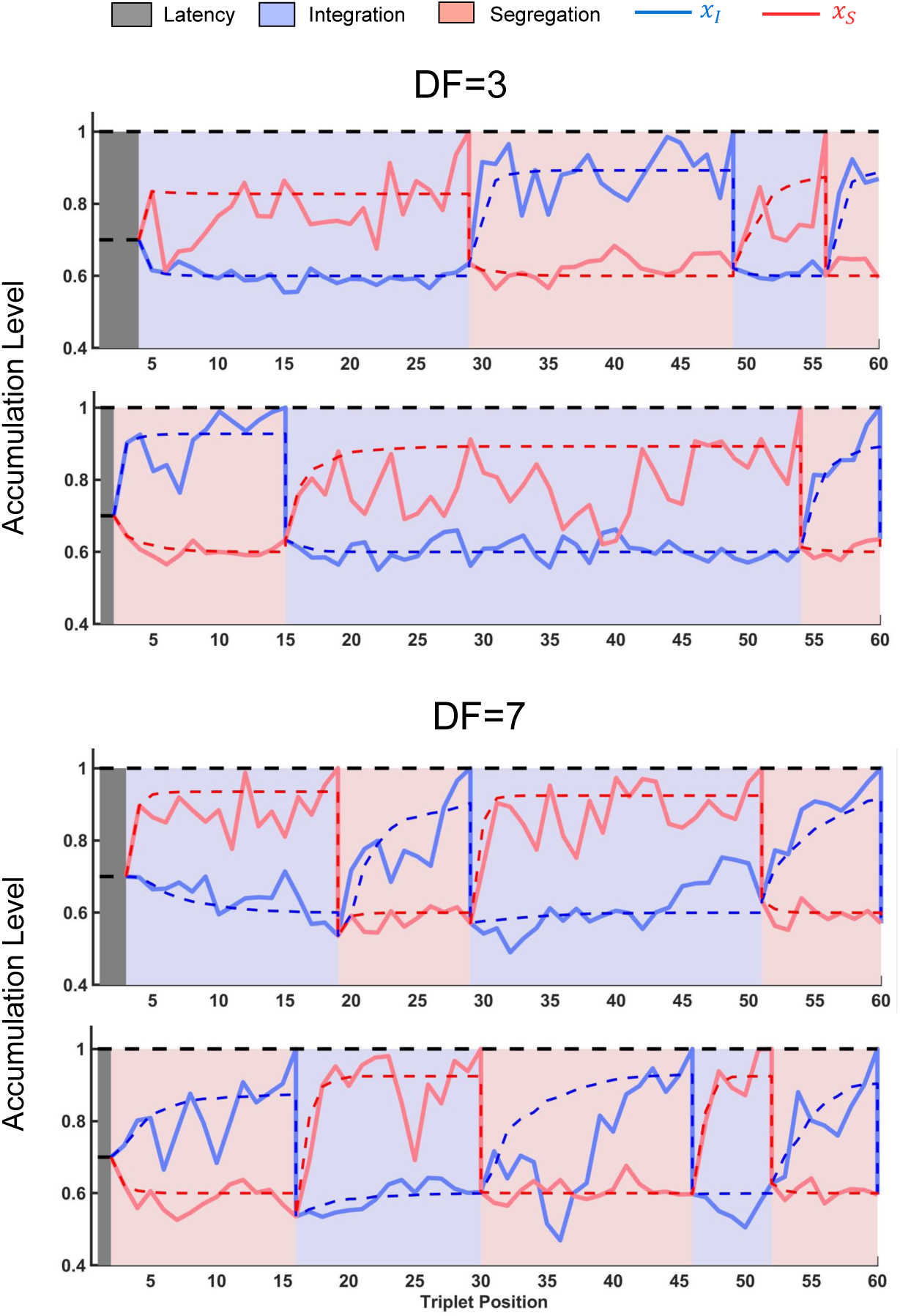
Exemplar time courses of accumulators in the EVA model, shown for *DF* =3 (top) and *DF* =7 (bottom). In only a few trials, 103 out of 675 for *DF* =3 and 220 out of 675 for *DF* =7, the first percept is segregation (see panels 2,4). During a cycle, the suppressed unit accumulates evidence against the current percept until it reaches the switching threshold. Then, a perceptual switch occurs and accumulators are reset to the same value. In the noise free case, the accumulators stabilize to their corresponding target values and there are no alternations. Such trajectories are depicted by dashed lines.

**S3 Fig.**
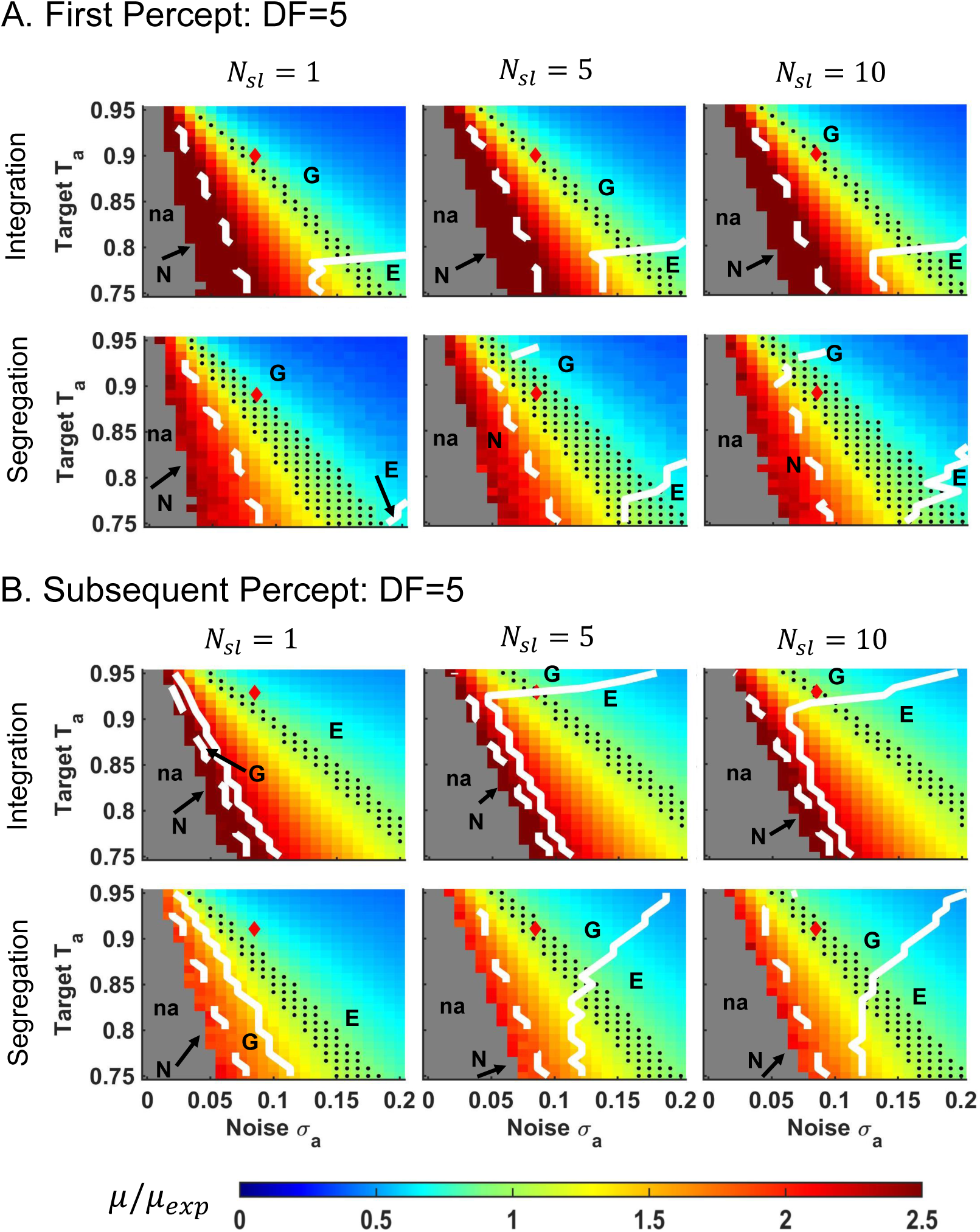
Mean percept durations (*µ*) and shape parameter values (*α*) in the EVA model are largely unaffected by changes in *N*_*sl*_, the number of units in the sampler layer. Some important differences occur, however, at *N*_*sl*_ = 1 (e.g. for subsequent durations). A two-parameter response diagram of the dependence of *µ* and *α* on target-against *T*_*a*_ and noise strength *σ*_*a*_ is shown for *DF* =5 and varying *N*_*sl*_ for A: first percepts and B: subsequent percepts. All parameters are chosen as described in Methods, except for *N*_*sl*_ (here *N*_*sl*_ = 1, 5 or 10). For comparison, see Fig 7, middle column; *N*_*sl*_ = 20 at *DF* =5. Red diamonds correspond to same parameter choices as in Fig 7 for *N*_*sl*_ = 20, as well. The heat map represents the ratio *µ/µ*_*exp*_ between model-generated *µ* and mean duration *µ*_*exp*_ from the behavioral data. Regions of no alternations (na) are colored in gray. Mean durations are much longer than their experimental counterparts *µ*_*exp*_ (region in warm colors), much shorter than *µ*_*exp*_ (region in cool colors), or close to *µ*_*exp*_ (within one standard error to *µ*_*exp*_; in green; black dots depict a discrete selection of values in the green region). The distributions of normalized percept durations are characterized by three distinct regions: for small *σ*_*a*_ the distributions are normal (region N, to the left of dashed-white line; *α* ≫ 3); for large *σ*_*a*_ the distributions are exponential (region E, to the right of solid-white line; *α* near 1); for intermediate values *σ*_*a*_ the distributions are gamma-like with shape close to that found experimentally (region G, between white contours; *α* ≈ 2 for first percepts and *α* ≈ 2.6 for subsequent percepts; *α* differs from *α*_*exp*_ by relative error up to 20% except for integration at *DF* =7 where it is up to 30%). As in Fig 7, middle column, the intersection of white contours with the sheet of black dots identifies parameter values that yield well-fit data. Note that at *N*_*sl*_ = 1 this intersection is empty for both subsequent percepts *I* and *S* (panel B, first column).

**S4 Fig.**
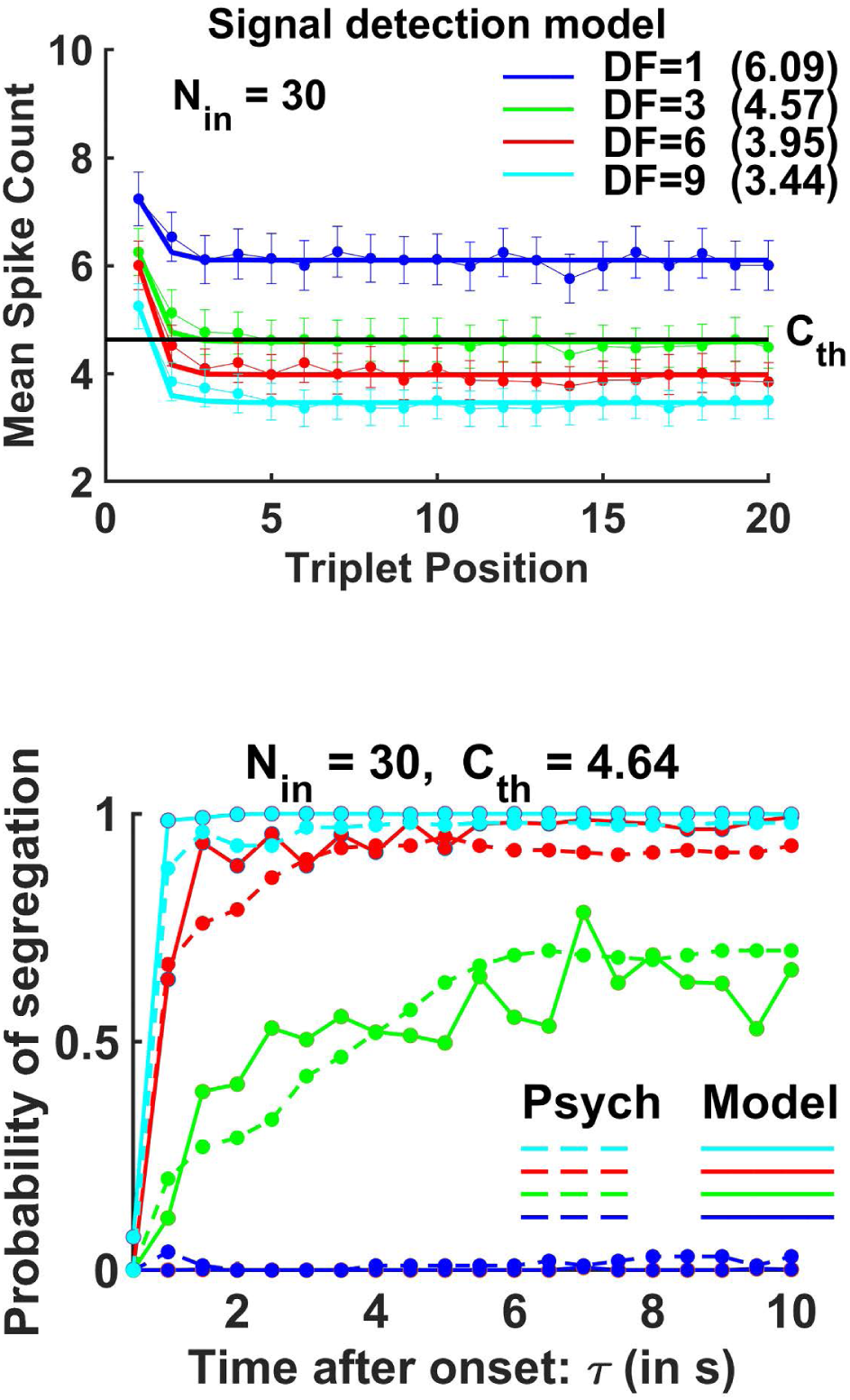
The signal detection algorithm for constructing a neurometric function (the probability of segregation as a function of time) generates acceptable buildup fits at *DF* = 1, 3, 6, 9. For comparison, see Micheyl et al (2005) [1]. Upper panel: mean spike counts *m*_*t*;*DF*_ (scatter points) at *A*-tone selective neurons in A1 during tone *B* were extracted from [1, (Fig.3A)]. They correspond to conditions *DF* =1 (blue), 3 (green), 6 (red), 9 (cyan), based on 10 s (20 triplets) long trials. The mean spike counts decrease exponentially and stabilize within a few seconds (solid curves for the exponential fits). The algorithm generates spike counts during *B*-tone by using Poisson processes of means *m*_*t*;*DF*_, and then average them over *N*_*in*_ neuronal units. The average values of the mean spike counts, including asymptotic values (written in parenthesis) at each *DF*, and the standard error to the mean (SEM) are computed over 675 trials. Lower panel: The signal detection algorithm constructs neurometric functions using numerical data from all *N*_*in*_ neuronal units. Parameters *N*_*in*_ and *C*_*th*_ are chosen to yield SEM similar to those observed in the spike count data [1, (Fig.3A)] and to yield the least-squares error of the experimental buildups (dashed, extracted from [1, (Fig.4)] and the computer-simulated neurometric functions (solid) for *DF* = 1, 3, 6, 9. The best approximation is obtained for *N*_*in*_ = 30, *C*_*th*_ = 4.64. Note: Statistics of percept durations were not reported in [1]; this prevented us from comparing these aspects of behavioral data from [1] to our numerically-generated duration distributions at *DF* =1, 3, 6, 9.

**S5 Fig.**
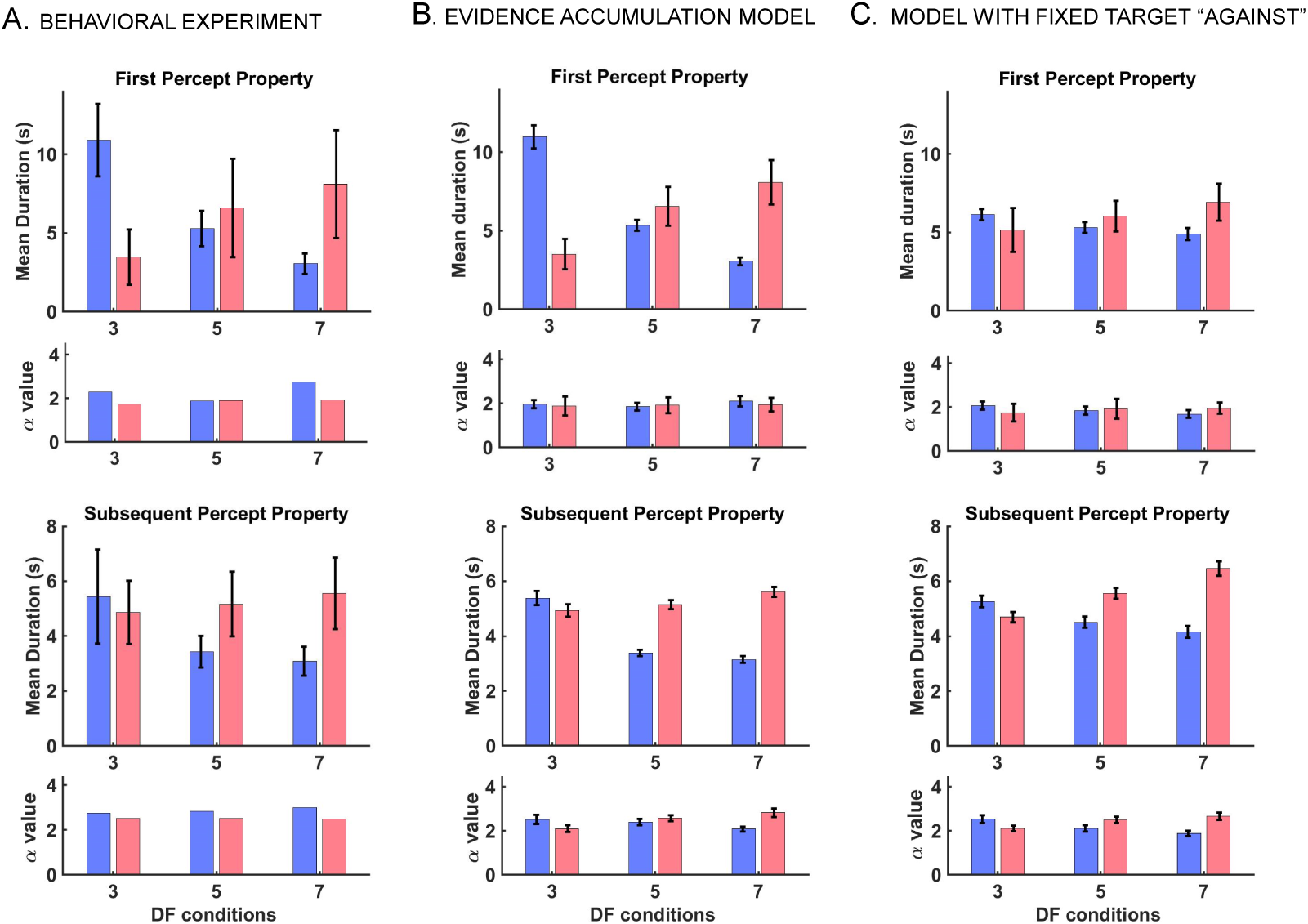
EVA model with fixed target-against value *T*_*a*_ across all conditions and percept types captures some, but not all, characteristics of perceptual alternations. For comparison, mean durations and shape parameter *α* of gamma distributions are shown for A: Experimental data; B: EVA model with optimized values for target-against (see Methods, Parameter values used in model simulations). EVA-generated results are identical to those in Figs 5 and 6; and C: Non-optimized EVA simulated with *T*_*a*_ = 0.9 across all *DF* = 3, 5, 7 and first and subsequent *I, S*. All other parameters are as in panel B. The mean durations from simulations follow the trend of experimental data which is decreasing/increasing with *DF* for *I*/*S* respectively. However, they fail to approximate well the entire set of behavioral data (e.g. approximations of mean first durations at *DF* =3 and *DF* =7 are inaccurate). On the other hand, gamma-fit shape values *α* are comparable to those from panels A and B. This is not surprising given that *α* depends mostly on the noise-level *σ*_*a*_, as shown in Fig 7.

